# Co-operation, competition and crowding: a discrete framework linking Allee kinetics, nonlinear diffusion, shocks and sharp-fronted travelling waves

**DOI:** 10.1101/077743

**Authors:** Stuart T. Johnston, Ruth E. Baker, D.L. Sean McElwain, Matthew J. Simpson

## Abstract

Invasion processes are ubiquitous throughout cell biology and ecology. During invasion, individuals can become isolated from the bulk population and behave differently. We present a discrete, exclusion-based description of the birth, death and movement of individuals. The model distinguishes between individuals that are part of, or are isolated from, the bulk population by imposing different rates of birth, death and movement. This enables the simulation of various co-operative or competitive mechanisms, where there is either a positive or negative benefit associated with being part of the bulk population, respectively. The mean-field approximation of the discrete process gives rise to 22 different classes of partial differential equation, which can include Allee kinetics and nonlinear diffusion. Here we examine the ability of each class of partial differential equation to support travelling wave solutions and interpret the long time behaviour in terms of the individual-level parameters. For the first time we show that the strong Allee effect and nonlinear diffusion can result in shock-fronted travelling waves. We also demonstrate how differences in group and individual motility rates can influence the persistence of a population and provide conditions for the successful invasion of a population.

## Introduction

Processes where individuals invade, and subsequently colonise, a region of space are prevalent in cell biology and ecology [1–11]. In cell biology, wound healing involves the invasion of fibroblasts into the wound space for tissue regeneration [7]. The invasion of glioma cells throughout the brain can lead to the formation of malignant brain tumours (glioblastoma) [1, 3, 10]. In ecology, the introduction and subsequent invasion of an alien species is a significant factor contributing to the extinction of native species [2, 11].

During invasion, individuals that become separated from the bulk population have been observed to have different behaviours to individuals within the bulk population [8, 12–15]. This is intuitive in ecological processes, as a decrease in the number of individuals within the bulk population can reduce the number of potential mates [13, 15–17] or lessen the efficacy of predator avoidance [14–16]. In cell biology, individual micrometastases have been observed to have reduced growth rates below a threshold size, which suggests that the presence of additional cells enhances the birth rate [12].

Continuum mathematical models of invasion processes have been studied extensively since the Fisher-Kolmogorov model was first proposed in 1937 [15, 18–30]. The Fisher-Kolmogorov model is a partial differential equation (PDE) description of the evolution of population density, where the temporal change in population density is attributed to a combination of linear diffusion and logistic growth [24, 27]. The Fisher-Kolmogorov model has been applied to various problems in cell biology and ecology [31–34]. The logistic growth term implies that the population density will always tend toward the carrying capacity [29]. This prediction does not reflect the observation that isolated individuals can experience a reduction in their birth rate [14]. This effect, known as the Allee effect, has two known forms. First, the strong Allee effect, where the growth rate is negative for sufficiently low densities [15, 35]. Second, the weak Allee effect, where the growth rate is reduced, but remains positive, at low densities [15]. Reaction-diffusion PDEs incorporating linear diffusion and Allee growth kinetics have been proposed and analysed [15, 19, 22, 23, 25, 26, 28, 30]. A key feature of interest for models of invasion is whether the PDE supports a travelling wave solution, where a wave front of constant shape moves through space with a constant speed. The sign of the wave speed indicates whether successful invasion occurs, and the magnitude of the wave speed provides an estimate of how quickly a population invades or recedes. More complicated descriptions of invasion processes with either Fisher or Allee kinetics and density-dependent nonlinear diffusion have been proposed, with the motivation of describing spatial aggregation or segregation [36–41].

A key feature of the Fisher-Kolmogorov model, and many extensions thereof, is that these PDE models are typically derived using phenomenologically-based arguments without incorporating information from an underlying stochastic description of individual-level behaviours. In this work we consider a relatively straightforward lattice-based discrete birth-death-movement model. An important characteristic of the model is that it explicitly accounts for crowding effects by only allowing for one agent per lattice site. Additionally, the rates at which birth, death and movement events occur depend on whether an agent is part of a group of agents or is isolated. We demonstrate that the standard continuum approximation of this discrete model can lead to either logistic or Allee kinetics, in an appropriate parameter regime. Furthermore, we demonstrate that imposing a different motility rate for agents that are isolated, compared to other agents, leads to a variety of density-dependent nonlinear diffusion functions. Previous studies have examined many different types of phenomenologically-based PDEs that are motivated in an *ad hoc* fashion. In contrast, our PDE description arises from a single, relatively simple, physically-motivated model. In Table 1 we highlight this generality, as the single discrete model gives rise to 22 different classes of PDE that describe the population-level behaviour.

While several of these PDEs have been studied previously, for completeness we examine the ability of each class of PDE to support travelling wave solutions. For certain classes of PDE, we present details of the travelling wave solutions for the first time. Interestingly, we obtain travelling wave solutions for PDEs that have nonlinear diffusivity functions with regions of negative diffusivity. Furthermore, we show that the strong Allee effect combined with these diffusivity functions can lead to novel shock-fronted travelling wave solutions. As these diffusivity functions are obtained directly from a discrete model, we can determine which competitive/co-operative individual-level mechanisms result in shock-fronted travelling wave solutions. Similarly, we are able to interpret the influence of motility on the persistence of a population, and highlight how this influence varies nonlinearly with the carrying capacity density. More generally, we provide new insight into the long time behaviour of an invasive population in terms of its individual-level properties.

**Table 1.** **Different classes of PDE associated with the discrete model in appropriate parameter regimes.** An Allee source term can correspond to either the weak, strong or reverse Allee effect. Degenerate diffusivity refers to the case where *F*(*C*\*)=*R*(*C*\*)=0 for some value *C*\*. A detailed analysis of all cases is presented in the Supplementary Material.

|  | Nonlinear<br>diffusivity | Degenerate<br>diffusivity | Number of<br>non-degenerate zeroes | Negative<br>diffusivity | Source<br>Term | Grouped<br>agent death | Relevant<br>previous analysis |
| --- | --- | --- | --- | --- | --- | --- | --- |
| Case 1 | x | x | 0 | x | Fisher | x | [18–21, 24, 25, 27, 29] |
| Case 2: Sub-case 1 | ✓ | x | 0 | x | Fisher | x | [42–44] |
| Case 2: Sub-case 2 | ✓ | ✓ | 0 | x | Fisher | x | [40, 45–47] |
| Case 2: Sub-case 3 | ✓ | x | 2 | ✓ | Fisher | x | [36] |
| Case 2: Sub-case 4 | ✓ | ✓ | 1 | ✓ | Fisher | x | [38] |
| Case 3 | x | x | 0 | x | Fisher | ✓ | [18–21, 24, 25, 27] |
| Case 4: Sub-case 1 | ✓ | x | 0 | x | Fisher | ✓ | [42–44] |
| Case 4: Sub-case 2 | ✓ | ✓ | 0 | x | Fisher | ✓ | [40, 45–47] |
| Case 4: Sub-case 3 | ✓ | x | 2 | ✓ | Fisher | ✓ | [36] |
| Case 4: Sub-case 4 | ✓ | ✓ | 1 | ✓ | Fisher | ✓ | [38] |
| Case 4: Sub-case 5 | ✓ | x | 1 | ✓ | Fisher | ✓ | [38] |
| Case 5 | x | x | 0 | x | Allee | x | [15, 19, 22, 23, 25, 26, 28, 30] |
| Case 6: Sub-case 1 | ✓ | x | 0 | x | Allee | x | [45, 48] |
| Case 6: Sub-case 2 | ✓ | ✓ | 0 | x | Allee | x | [41, 49] |
| Case 6: Sub-case 3 | ✓ | x | 2 | ✓ | Allee | x | [37] |
| Case 6: Sub-case 4 | ✓ | ✓ | 1 | ✓ | Allee | x | [39] |
| Case 7 | x | x | 0 | x | Allee | ✓ | [15, 19, 22, 23, 25, 26, 28, 30] |
| Case 8: Sub-case 1 | ✓ | x | 0 | x | Allee | ✓ | [45, 48] |
| Case 8: Sub-case 2 | ✓ | ✓ | 0 | x | Allee | ✓ | [41, 49] |
| Case 8: Sub-case 3 | ✓ | x | 2 | ✓ | Allee | ✓ | [37] |
| Case 8: Sub-case 4 | ✓ | ✓ | 1 | ✓ | Allee | ✓ | [39] |
| Case 8: Sub-case 5 | ✓ | x | 1 | ✓ | Allee | ✓ | [39] |

## Results

We consider a discrete lattice-based exclusion process where agents undergo birth, death and movement events. We distinguish between isolated agents and grouped agents by imposing different rates of birth, death and movement depending on whether an agent has zero or at least one nearest-neighbour agent, respectively. A more detailed description of the discrete model is presented in the Methods. To derive a continuum limit PDE description of the discrete model [58, 63] we consider the change in occupancy of a lattice site *j* during a single time step of duration *τ*, and obtain

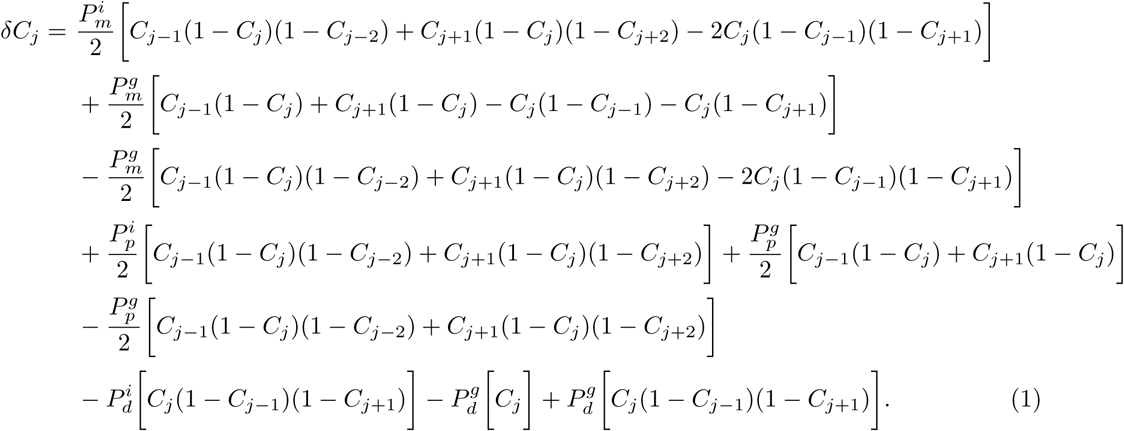

Here, *C_j_* represents the probability that the site *j* is occupied and, therefore, 1 – *C_j_* represents the probability that the site is vacant [63]. Furthermore, as products of probabilities are interpreted as net transition probabilities, the usual assumption that the occupancy of lattice sites are independent is made [58, 61, 64–66].

Note that *C_j_* is the total occupancy of site *j*, that is, the sum of the occupancy of isolated agents and the occupancy of grouped agents at that site. We now interpret the terms on the right-hand side of Equation (1) in terms of the physical change in lattice occupancy. The positive terms proportional to 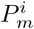 correspond to isolated agents moving into site *j*, while the negative terms correspond to isolated agents moving out of site *j*. Each term consists of three factors. For the negative terms, these factors are the probability that site *j* is occupied, and the probabilities that sites *j* – 1 and *j* + 1 are vacant. For the positive terms, the three factors are the probability that site *j* ± 1 is occupied, and the probabilities that sites *j* and *j* ± 2 are vacant. The third factor is required to ensure that the term describes isolated agents. The positive/negative terms proportional to the first 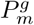 term on the right-hand side of Equation (1) correspond to grouped agents moving in/out of site *j*. These terms consist of two factors; the probability that the selected site is occupied and the probability that the target site is vacant. The second 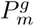 term ensures that the isolated agents are not counted twice. The remaining terms can be interpreted similarly; products of probabilities that specific sites are occupied or vacant that describe the change of occupancy of a site in response to a birth or death event.

To obtain a PDE description we divide Equation (1) by *τ* and consider *C_j_* as a continuous function, *C*(*x*, *t*). We expand *C*(*x*, *t*) in a Taylor series around *x* = *j*∆, truncating terms of 𝒪(∆^3^), where ∆ is the lattice spacing [58, 63]. Taking the limit ∆ → 0 and *τ* → 0 such that ∆^2^/*τ* is held constant [60, 63, 67] gives

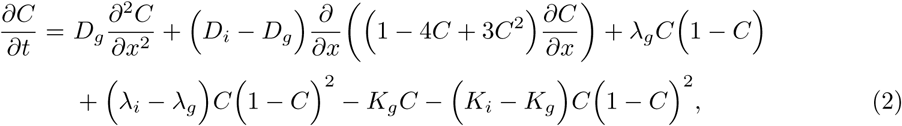

where

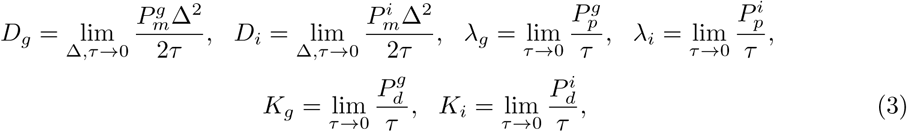

with the further assumption that 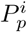, 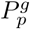, 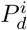, 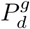 are 𝒪(*τ*) [63]. The individual-level parameters are treated as being interchangeable with the continuum-level parameters as defined in (3). All implementations of the discrete model in this work have ∆ = *τ* = 1.

It is convenient to write Equation (2) in conservation form

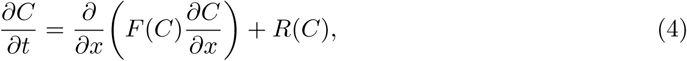

where

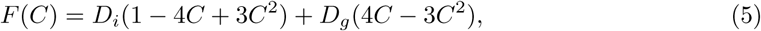

is the nonlinear diffusivity function, and

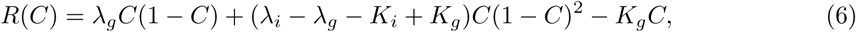

is the source/sink term.

The aims of this work are to first illustrate that the very different types of behaviour encoded in the discrete model are also reflected in the solution of Equation (2). Once we have demonstrated this connection, we focus on examining travelling wave solutions for the 22 different classes embedded within Equation (2), as summarised in Table 1. In the main document we highlight novel and key results for specific classes of PDEs resulting from the discrete model, and provide relevant discussion about the implications of the long time population behaviour. A more thorough investigation of the travelling wave solutions arising from all 22 classes of PDEs is presented in the Supplementary Material.

Twenty identically-prepared realisations of the discrete model are presented in Figures 1(a)-(f) for two different parameter regimes. In the first parameter regime, where 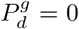, the initially-occupied region of the lattice remains fully occupied, as shown in Figures 1(a)-(c). When we introduce 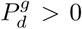, as shown in Figures 1(d)-(f), the initially-occupied region of the lattice becomes partially vacant as time increases. We also compare the average discrete behaviour and the corresponding numerical solution of Equation (2) in Figures 1(g)-(h). This comparison shows that the solution of the continuum PDE matches the average discrete behaviour well, and predicts both the spread of the agent population in Figure 1(g) and the decrease in agent density in Figure 1(h).

The governing PDE, Equation (2), can be simplified in specific parameter regimes. While several of these simplified PDEs have been studied extensively, we summarise all non-trivial cases for completeness. It is instructive to consider each case and discuss the implications of the long term behaviour in terms of the discrete model parameters, as previous derivations of these PDEs have arisen from a variety of *ad hoc* arguments rather than working with a single unifying model. In Table 1 we summarise the salient features of 22 different special cases of Equation (2). The nonlinear diffusivity function, *F*(*C*), has four key properties:

- *F*(*C*) can either be a constant, or a function of the density of individuals;
- *F*(*C*) can be degenerate, which implies that at one or more densities, *C*\*, we have *F*(*C*\*) = *R*(*C*\*) = 0;
- *F*(*C*) can be zero at values of *C*\* that are non-degenerate, that is, *F*(*C*\*) = 0, *R*(*C*\*) ≠ 0. In our model, this can occur at either zero, one or two different values of *C*; and
- *F*(*C*) can be negative for an interval of *C* values.

**Figure 1.**
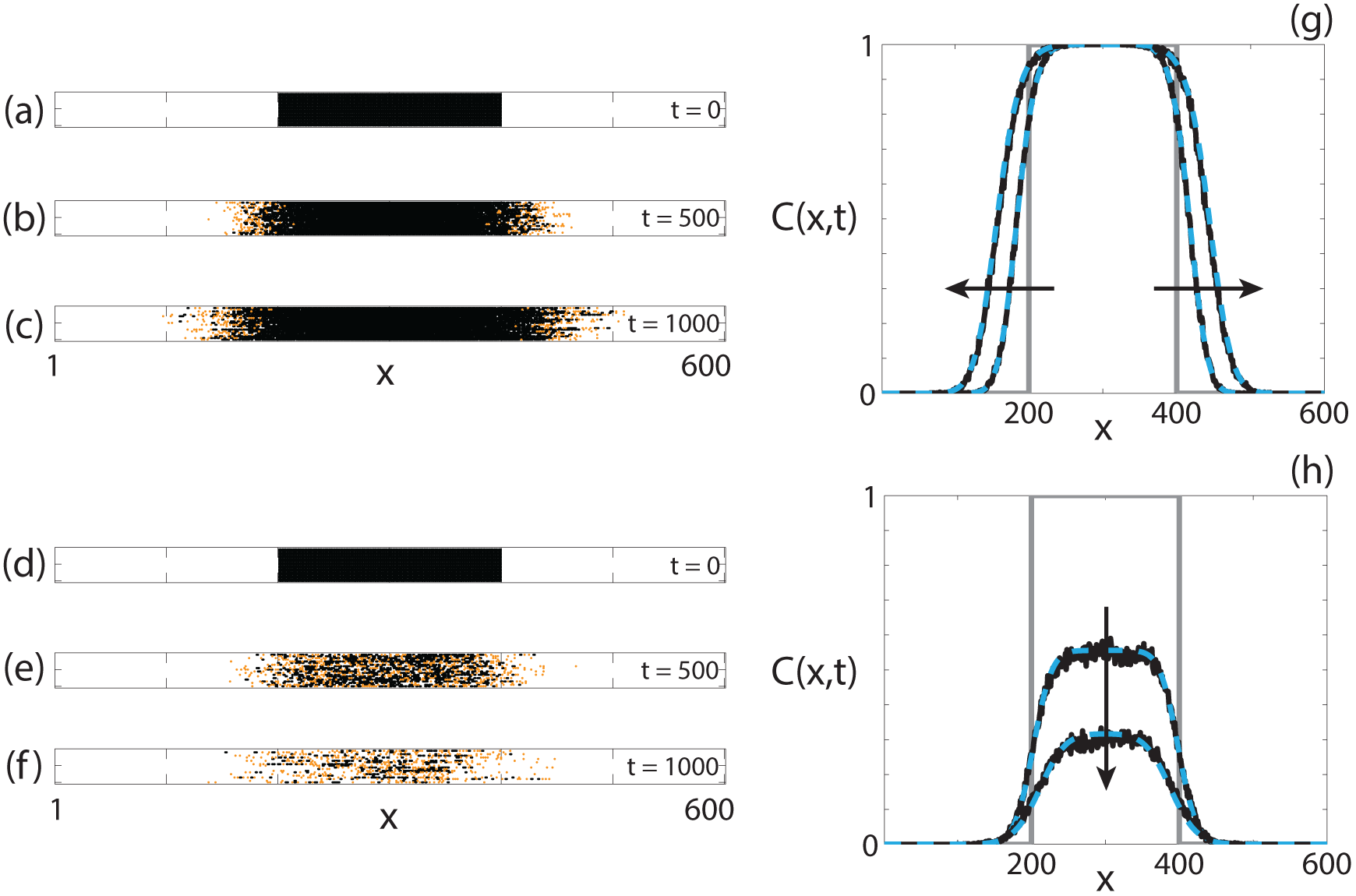
**Comparison of the discrete model and the continuum approximation.** (a)-(f) 20 identically-prepared realisations of the discrete model at (a), (d) *t* = 0; (b), (e) *t* = 500; (c), (f) *t* = 1000. The discrete model simulations correspond to (a)-(c) 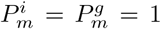, 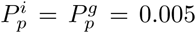, 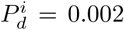, 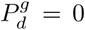; (d)-(f) 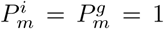, 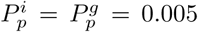, 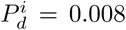, 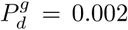. For all simulations *τ* = 1, ∆ = 1. (g)-(h) Comparisons between the averaged discrete model (black, solid) and the numerical solution of Equation (2) (cyan, dashed) at *t* = 0, *t* = 500 and *t* = 1000 for the parameters in (a)-(c) and (d)-(f), respectively. The grey lines indicate the initial condition and the arrow indicates the direction of increasing time. For all discrete solutions, *M* = 1000, *X* = 600, ∆ = *τ* = 1. For all continuum solutions, *δx* = 1, *δt* = 0.1, ∊ = 10^−6^.

The source term, *R*(*C*), has two key properties:

- *R*(*C*) can represent either Fisher kinetics (logistic growth) or Allee kinetics (bistable); and
- the grouped agent death rate, 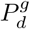, can be zero or non-zero. If the rate is non-zero, the carrying capacity density is reduced.

There are three different types of Allee kinetics considered in this work; weak, strong and reverse. We consider these three kinetics together for brevity, as it is relatively simple to change the parameter regime to alter the type of Allee effect without changing the competitive/co-operative mechanism described. The reverse Allee effect, which we describe here for the first time, refers to a growth rate that is reduced at high density, compared to logistic kinetics, but remains positive.

### Fisher kinetics

The choice of whether the birth and death mechanisms imposed in the discrete model are neutral or are competitive/co-operative determines the form of the source term. If both 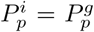 and 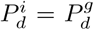, and hence both the birth and death mechanisms are neutral, the source term represents Fisher kinetics and Equation (2) simplifies to

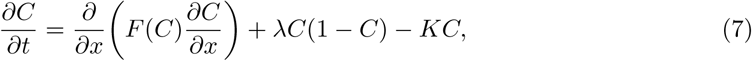

where *λ* = *λ*_*i*_ = *λ*_*g*_ and *K* = *K_i_* = *K*_*g*_. Transforming Equation (7) into travelling wave coordinates *z* = *x* – *υt*, where *υ* is a constant wave speed and −∞ < *z* < ∞, results in

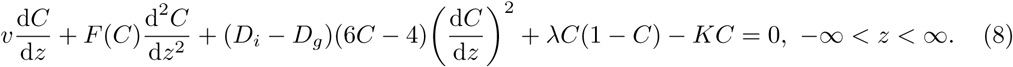

Substituting *U* = d*C*/d*z* allows Equation (8) to be expressed as a system of ordinary differential equations (ODEs)

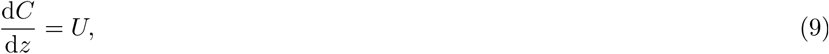

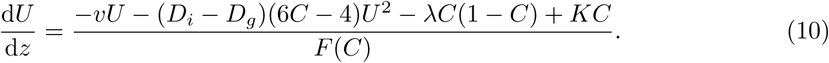

The equilibrium points of Equations (9)–(10) occur at (*C*, *U*) = (0,0) and (*C*, *U*) = (*S*, 0), where *S* = (*λ* – *K*)/*λ*. The range of physically relevant *C* values correspond to 0 ≤ *C* ≤ *S*. Hence the carrying capacity density, S, determines the numbers of times that *F*(*C*) = 0 for physically relevant *C* values. As such, we introduce a new variable *C̄* = *C/S* such that the agent density is scaled by the carrying capacity and the zeroes of *R*(*C̄*) occur at *C̄* = 0 and *C̄* = 1.

**Figure 2.**
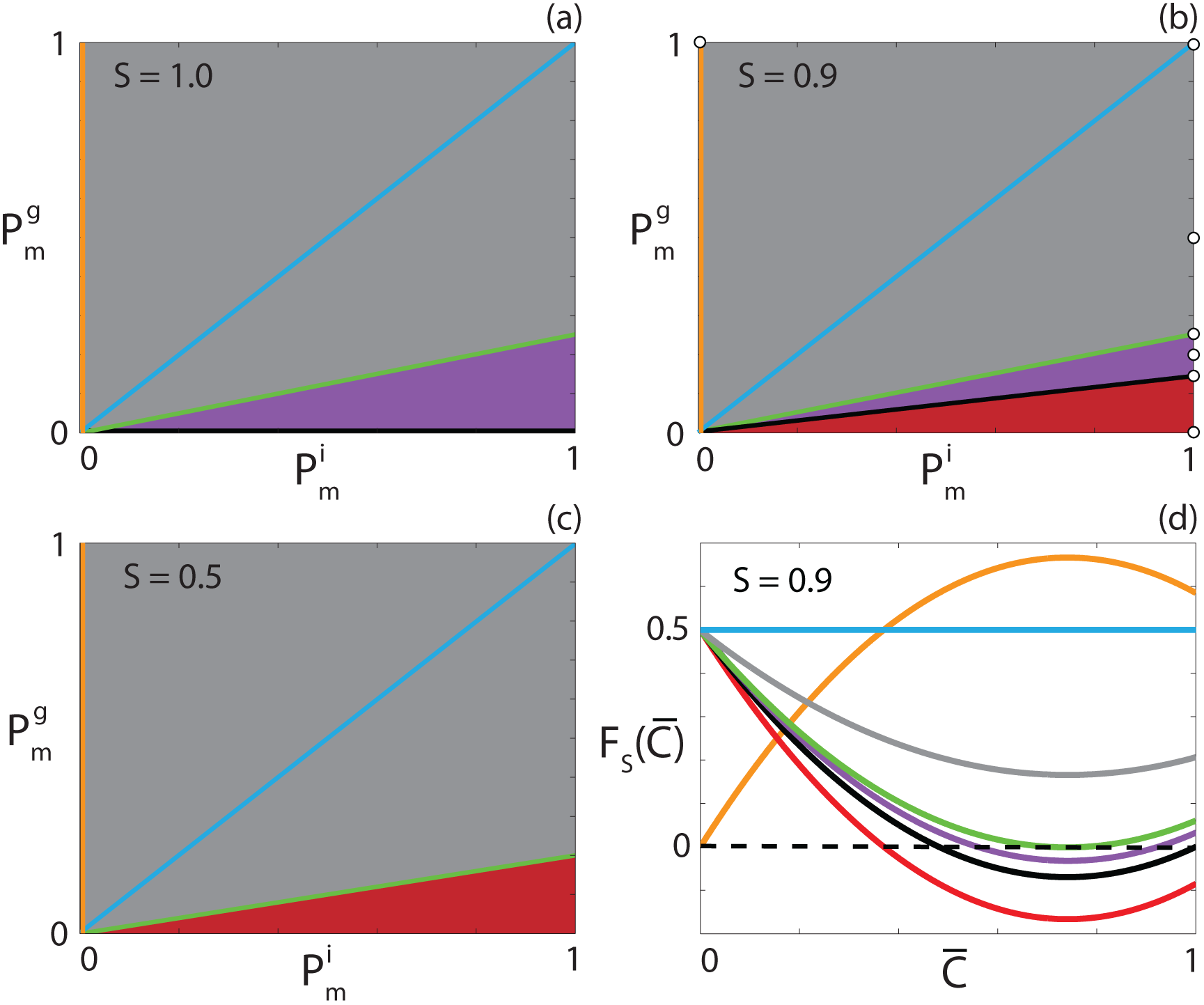
**Classification of** *F_s_*(*C̄*) for different carrying capacity densities. (a)-(c) Type of *F_s_*(*C̄*) function for 0 ≤ *C̄* ≤ 1 for the parameter space 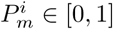 and 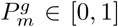 with (a) *S* = 1.0; (b) *S* = 0.9; (c) *S* = 0.5. Grey regions correspond to parameter pairs that result in strictly positive *F_s_*(*C̄*), purple regions correspond to parameter pairs that result in positive-negative-positive *F_s_*(*C̄*) and red regions correspond to parameter pairs that result in positive-negative *F_s_*(*C̄*). Cyan, orange and black lines correspond to constant, extinction-degenerate non-negative and capacity-degenerate positive-negative *F_s_*(*C̄*) curves, respectively. (d) Example *F_s_*(*C̄*) for each region in (b). The white circles in (b) denote the parameter pairs used to generate the curves in (d).

Transforming Equation (7) in terms of (*C̄*), we obtain

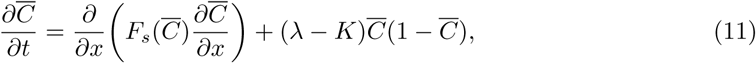

where *F_s_*(*C̄*) = *F*(*SC̄*) = *D_i_*(1 – 4*SC̄* + 3(*SC̄*)^2^) + *D_g_*(4*SC̄* – 3(*SC̄*)^2^). Equation (11) is the Fisher-Kolmogorov equation in terms of *C̄* with a nonlinear diffusivity function, *F_s_*(*C̄*). This new nonlinear diffusivity function has different properties depending on *S*, *D_i_* and *D_g_*. To highlight this, Figure 2 shows the 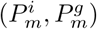 parameter space for three different choices of *S* and the qualitative behaviour of the corresponding *F_s_*(*C̄*) function. For all *S* values, parameter pairs that result in a constant *F_s_*(*C̄*) are highlighted in cyan. All parameter pairs that result in *F_s_*(*C̄*) > 0 for 0 ≤ *C̄* ≤ 1 are denoted by the grey regions. This type of diffusivity function is referred to as *strictly positive*. Similarly, for all *S* values, there are parameter pairs that result in *F_s_*(0̄) = 0, and *F_s_*(*C̄*) > 0 otherwise, which are highlighted in orange. We refer to this type of diffusivity function as *extinction-degenerate non-negative*.

For *S* = 1, presented in Figure 2(a), 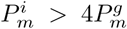, denoted in purple, results in an interval *α* < *C̄* < *β*, *α* < *β* < 1, where *F_s_*(*C̄*) < 0. We refer to this type of nonlinear diffusivity function *as positive-negative-positive*. Decreasing *S* to 0.9, presented in Figure 2(b), we observe that the purple region again occurs for 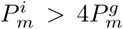. However, if 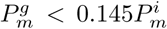, highlighted in red, *F_s_*(*C̄*) < 0 for *ω* < *C̄* ≤ 1, and hence *F*_*s*_(*C̄*) has only one zero in 0 ≤ *C̄* ≤ 1. This type of nonlinear diffusivity function is not observed with *S* = 1 and we refer to it as *positive-negative*. Specifically, this behaviour occurs when 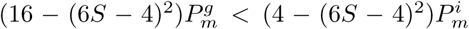 and 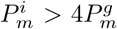. Furthermore, this implies that for *S* < 2/3 there are no 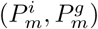 values that correspond to positive-negative-positive *F*_*s*_(*C̄*). A choice of 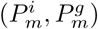 that demonstrates this is shown in Figure 2(c). Unlike in Figures 2(a)-(b), we see that there is no purple region. Finally, if 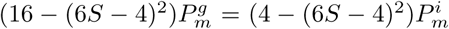, highlighted in black, *F_s_*(1) = 0 and *F*_*s*_(*C̄*) < 0 for *ω* < *C̄* < 1, which we refer to as *capacity-degenerate positive-negative*. Note that for *S* < 1/3, *F*_*s*_(*C̄*) ≥ 0 for 0 ≤ *C̄* ≤ 1. An example *F*_*s*_(*C̄*) curve for each type of diffusivity function is presented in Figure 2(d). PDE models that contain diffusivity functions with a region of negative diffusivity have been considered previously. However, these models either do not contain a source term or consider source terms that do not support travelling wave solutions [70–72]. Hence the model and analysis considered in this work is significantly different to the previous studies.

For all combinations of neutral, competitive and co-operative mechanisms that give rise to a reaction-diffusion equation with Fisher kinetics we examine the ability of the equation to give rise to long time travelling wave solutions. While details of the travelling wave solutions for certain types of diffusivity functions have been presented previously, we summarise the key features of the travelling wave solutions in tabular form for all cases for completeness in Table 2. For the cases where solution profiles have not been presented previously we provide more detailed discussion. A detailed analysis for each case is presented in the Supplementary Material.

**Table 2.** Classification of travelling wave solutions arising from different classes of PDEs with Fisher kinetics. Highlighted entries refer to cases analysed in detail in the manuscript; the other cases are analysed in the Supplementary Material.

| Diffusivity function classification | Travelling wave | Front type | Direction |
| --- | --- | --- | --- |
| Constant | Yes | Smooth | Positive |
| Strictly positive | Yes | Smooth | Positive |
| Extinction-degenerate non-negative | Yes | Sharp | Positive |
| Positive-negative-positive | Yes | Smooth | Positive |
| Capacity-degenerate positive-negative | Yes | Smooth | Positive |
| Positive-negative | Yes | Smooth | Positive |

#### Positive-negative-positive nonlinear diffusivity function

The first diffusivity function we examine in detail is the positive-negative-positive nonlinear diffusivity function, where *F*_*s*_(*C̄*) < 0 for an interval *α* < *C̄* < *β*. The simplest positive-negative-positive *F*_*s*_(*C̄*) occurs where 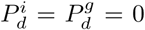 and hence *S* = 1. For these parameters, *F*_*s*_(*C̄*) = *F*(*C*). Note that introducing non-zero 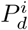 and 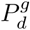 merely scales the governing equation and hence extending this analysis to cases with non-zero agent death is straightforward, provided that *F*_*s*_(*C̄*) has two zeroes on the interval 0 < *C̄* < 1. Parameters that result in a positive-negative-positive *F*_*s*_(*C̄*) are highlighted in purple in Figure 2 and, for this case, with 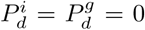, occur when 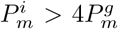. For positive-negative-positive *F*(*C*), Equation (10) is singular at *C* = *α* and *C* = *β*, where the interval of *F*(*C*) < 0 is given by

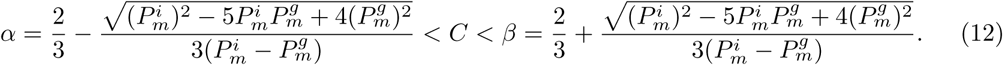

The singularities at *C* = *α* and *C* = *β* cannot be removed using a stretching transformation (Supplementary Material) since *R*(*α*) ≠ 0 and *R*(*β*) ≠ 0. However, it is possible for d*U*/d*z* to be finite at *C* = *α* and *C* = *β* if *U_α_* and *U_β_* exist such that

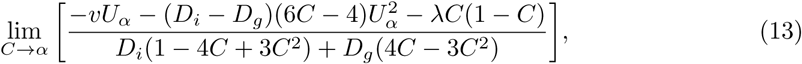

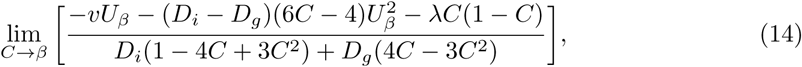

are both finite. This requires the numerator in the expressions (13)–(14) vanish at *C* = *α* and *C* = *β*, respectively. As such, *U_α_* and *U_β_* are obtained by solving the system

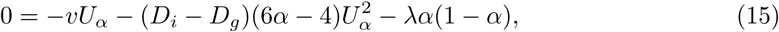

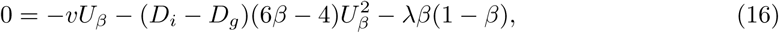

resulting in 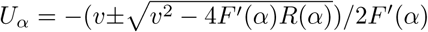 and 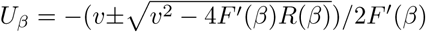. We note that as *R*(*C*) ≥ 0 for 0 ≤ *C* ≤ 1, and that *F'*(*α*) ≤ 0 for all possible *α* values, *U_α_* will be real-valued. Subsequently, we have a wave speed condition that 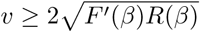, as *F'*(*β*) ≥ 0 for all possible *β* values. Ferracuti *et al*. [36] prove that the minimum wave speed, *υ*^*^, is greater than a threshold value, which in turn is greater than max{*R*'(0)*F*(0), *F*'(*β*)*R*(*β*)}. Therefore, *U_β_* will also always be real-valued.

Applying L'Hopital's Rule to Equation (10), we obtain

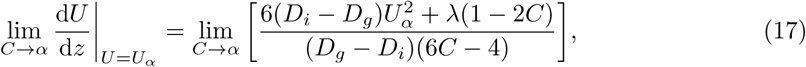

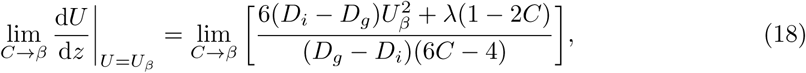

which are finite provided that *α* ≠ 2/3 and *β* ≠ 2/3. For the system of Equations (9)–(10), we have two straight lines in the phase plane where d*U*/d*z* is infinite, at *C* = *α* and *C* = *β*. These kind of lines have previously been called *walls of singularities* for hyperbolic models related to chemotactic and haptotactic invasion [52]. For a smooth solution trajectory joining the two equilibrium points on opposite sides of the wall of singularities, we require that the trajectory passes through the wall of singularities. This implies that the solution trajectory must pass through the wall of singularities at the special points, (*α*, *U_α_*) and (*β*, *U_β_*), known as *holes in the wall* [52, 53]. Otherwise, a smooth heteroclinic orbit between (1, 0) and (0, 0) cannot exist, as lim_*C*→*α*_ |*U*| →∞ and lim_*C*→*β*_ |*U*| →∞. As *U_α_* and *U_β_* are real valued and the limits in Equations (11)–(12) are finite, the holes in the wall always exist for Fisher kinetics.

We superimpose the numerical solution of Equation (7) in (*C*, *U*) co-ordinates on the phase plane for the system (9)–(10) in Figures 3(a) and 3(d). Details of the numerical techniques used to solve Equation (7) and to generate the phase planes are given in the Methods. The numerical solution appears to form a heteroclinic orbit between (1, 0) and (0, 0) in both cases, and passes through the holes in the wall of singularities, denoted using purple circles. Continuum models with negative diffusivity and no source terms have been relatively well studied, and exhibit shock behaviour across the region of negative diffusion [50, 51]. Interestingly, our solution does not include a shock and is instead smooth through the region of negative diffusion.

**Figure 3.**
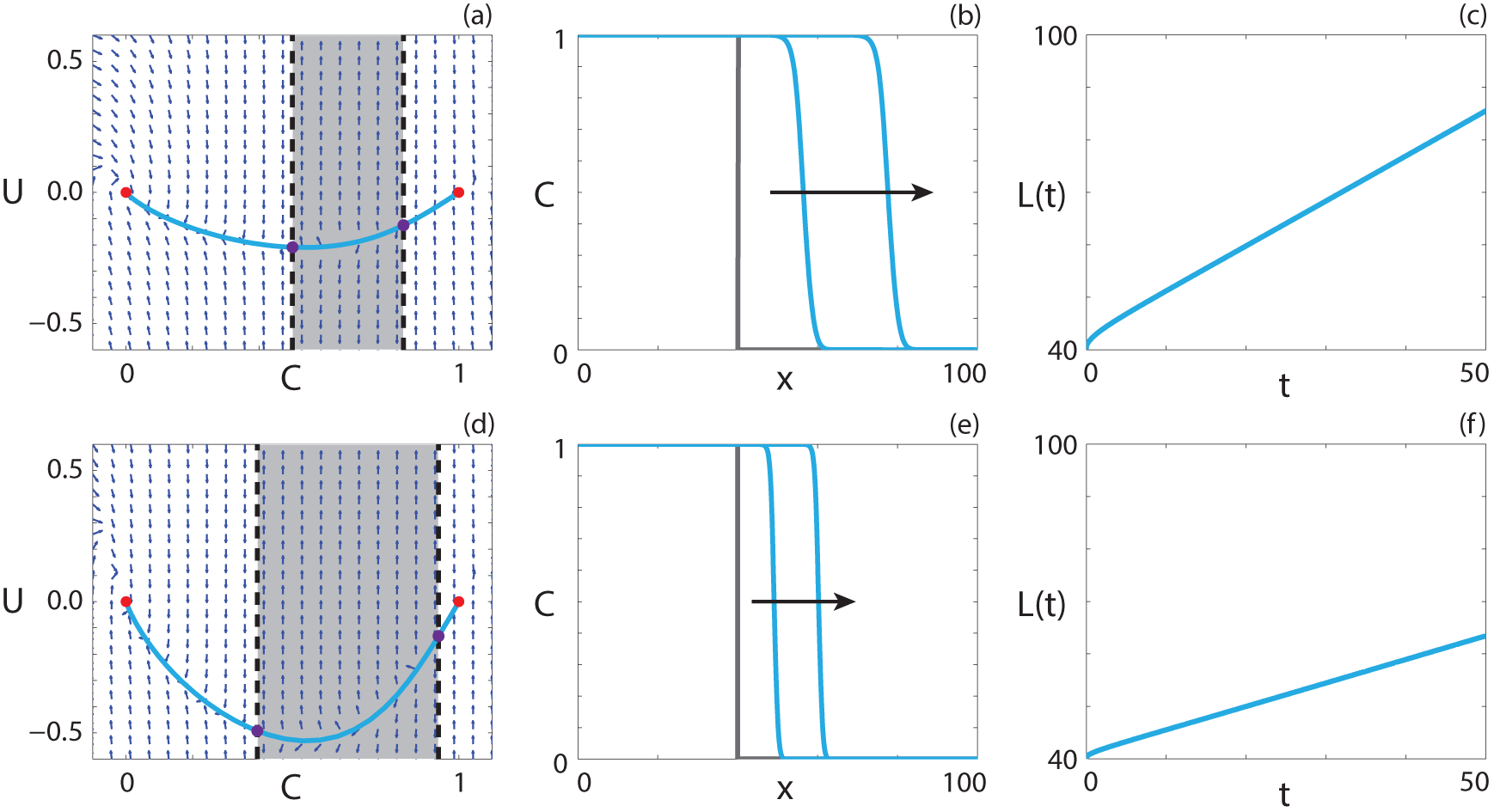
**Travelling wave behaviour for Equation (7) with positive-negative-positive *F*(*C*) (Case 2.3).** (a), (d) Phase plane for the system (9)–(10) with the numerical solution of Equation (7), in (*C*, *U*) co-ordinates, superimposed. The grey region corresponds to values of *C* where *F*(*C*) < 0. The dashed black lines denote a wall of singularities. Red circles correspond to equilibrium points and purple circles correspond to holes in the wall. (b), (e) Numerical solution of Equation (7) at *t* = 100 and *t* = 200. The grey lines indicate the initial condition and the arrows indicate the direction of increasing time. (c), (f) The time evolution of the position of the leading edge of the travelling wave solution, *L*(*t*). All results are obtained using 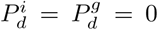, *δx* = 0.01, *δt* = 0.01, *∊* = 10^−6^ and (a)-(c) 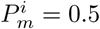, 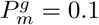, 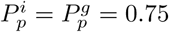, *υ* = 0.864; (d)-(f) 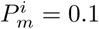, 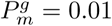, 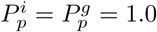, *υ* = 0.448.

The numerical solution of Equation (7) at *t* = 100 and *t* = 200 is shown in Figures 3(b) and 3(e), confirming that the waveform does not change with time. To quantify the wave speed we calculate the time evolution of the leading edge, *L*(*t*) = *x_f_* such that *C*(*x_f_*, *t*) ≈ 1 × 10^−4^. If the solution of Equation (7) forms a travelling wave, *L*(*t*) will tend to a straight line with slope *υ*, as *t*→∞ In Figures 3(c), and 3(f), we observe that *L*(*t*) is approximately linear with slope *υ*, and hence the solution of Equation (7) moves with approximately constant speed at long times. Overall, these features suggest that the solution of Equation (7) with positive-negative-positive *F*(*C*) approaches a travelling wave.

#### Capacity-degenerate positive-negative nonlinear diffusivity function

For the special case where 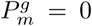, *F_s_*(1) = 0. Again, we consider the case with zero agent death for simplicity, and note that it is straightforward to extend the analysis for cases with non-zero agent death. As *F*(*C*) is degenerate at *C* = 1, it is intuitive to expect there could be sharp-fronted travelling wave solutions, with the sharp front near *C* = 1, similar to the results in [46] and in the Supplementary Material. However, unlike these cases, we have an interval 1/3 < *C* < 1 where *F*(*C*) < 0. To determine whether this negative diffusivity influences the presence of sharp fronts, we follow the approach of Maini *et al*. [38], who show that the existence of travelling waves for reaction-diffusion equations with capacity-degenerate positive-negative *F*(*C*) can be determined by considering the existence of travelling waves for

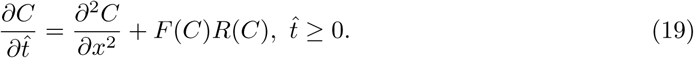

The restriction on *t̂* implies *F*(*C*) > 0. As *F*(*C*) < 0 for 1/3 < *C* < 1, Equation (19) is only equivalent to Equation (7) for 0 ≤ *C* ≤ 1/3. For 1/3 ≤ *C* ≤ 1, Equation (7) is equivalent to

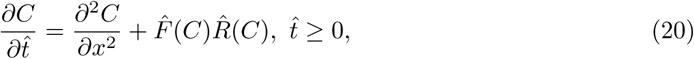

where *F̂*(*C*) = −*F*(1 – *C*) and *R̂*(*C*) = *R*(1 – C) [38]. Equations (19)–(20) have minimum travelling wave speeds 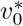 and 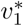. Maini *et al*. [38] prove that sharp fronts in the travelling wave near *C* = 1 only exist if *F*(1) = 0 and 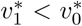. The first condition is obviously satisfied, while the second can be determined by considering the behaviour of the equivalent ordinary differential systems in travelling wave coordinates in the neighbourhood of the equilibrium points. Both equations have minimum wave speed conditions, 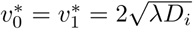, to obtain physically-relevant heteroclinic orbits, and hence travelling wave solutions with a sharp region near *C* = 1 do not exist.

Travelling wave behaviour for a parameter regime with *F*(1) = 0 is shown in Figure 4. The equilibrium point at (1, 0) is also a hole in the wall. The solution trajectory forms a heteroclinic orbit between (1, 0) and (0, 0), and passes through the region of *C* where *F*(*C*) < 0. Although *F*(1) = 0, we do not observe a solution trajectory corresponding to a sharp front, as for capacity-degenerate non-negative *F*(*C*) (Supplementary Material). This result is consistent with the analysis of Maini *et al*. [38]. The numerical solution of Equation (4), presented in Figure 4(b), has a relatively steep front but is not sharp near *C* = 1. As *L*(*t*), presented in Figure 4(c), becomes linear as *t* increases and the waveform in Figure 4(b) are consistent, the numerical solution of Equation (7) with *F*(1) = 0 appears to form a classic travelling wave.

**Figure 4.**
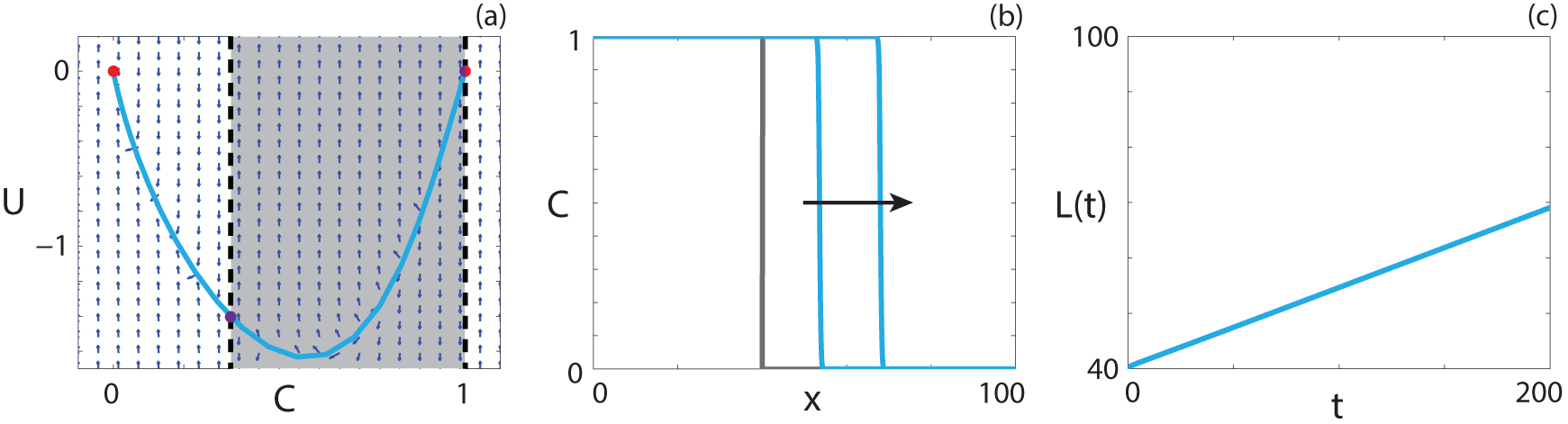
**Travelling wave behaviour for Equation (7) with capacity-degenerate positive-negative *F*(*C*) (Case 2.4).** (a) Phase plane for the system (9)–(10) with the numerical solution of Equation (7), in (*C*, *U*) co-ordinates, superimposed. The grey region corresponds to values of *C* where *F*(*C*) < 0. The dashed black lines denote two walls of singularities. Red circles correspond to equilibrium points and purple circles correspond to holes in the wall. (b) Numerical solution of Equation (7) at *t* = 100 and *t* = 200. The grey lines indicate the initial condition and the arrow indicates the direction of increasing time. (c) The time evolution of the position of the leading edge of the travelling wave solution, *L*(*t*). All results are obtained using 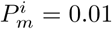, 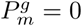, 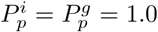, 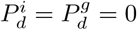, *δx* = 0.01, *δt* = 0.01, *∊* = 10^−6^, *υ* = 0.1433.

#### Positive-negative nonlinear diffusivity function

The positive-negative case, where *F*_*s*_(*C̄*) > 0 for 0 ≤ *C̄* < *ω* and *F*_*s*_(*C̄*) < 0 for *ω* < *C̄* ≤ 1, cannot occur with *K* = 0. It is instructive to examine whether stable travelling wave solutions of Equation (7) exist in such a case, as the non-zero equilibrium point now occurs in the region where *F*_*s*_(*C̄*) < 0. If we perform standard linear analysis on Equations (9)–(10), the Jacobian at (*S*, 0) has eigenvalues 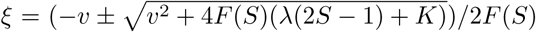, which implies that the equilibrium is an unstable node provided 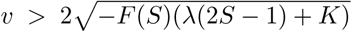. The negative sign is present as *F*(*S*) < 0 for positive-negative *F*_*s*_(*C̄*). The Jacobian at (0, 0) has eigenvalues 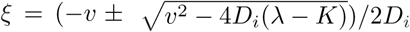 which is a stable node provided that 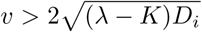. While there are infinitely many solution trajectories out of the unstable node, we require that the solution trajectory passes through the hole in the wall, and hence there is a single solution trajectory that forms a heteroclinic orbit.

**Figure 5.**
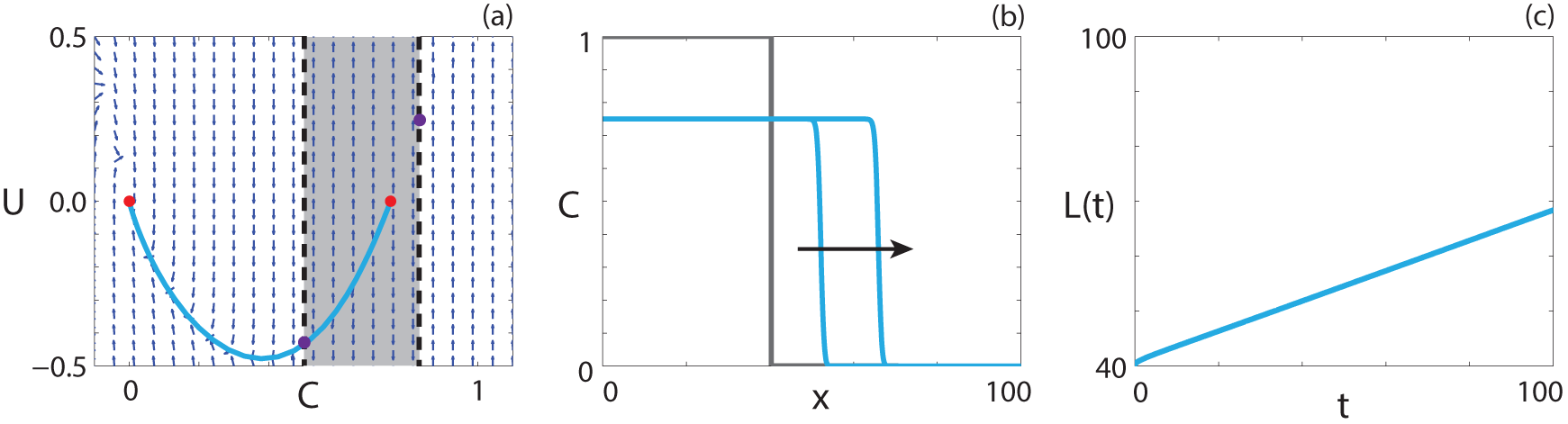
**Travelling wave behaviour for Equation (7) with positive-negative *F_s_*(*C̄*) (Case 4.5).** (a) Phase plane for the system (9)–(10) with the numerical solution of Equation (7), in (<u>*C,U*</u>) co-ordinates, superimposed. The grey region corresponds to values of *C̄* where *F*_*s*_(*C̄*) < 0. The dashed black lines denote a wall of singularities. Red circles correspond to equilibrium points and purple circles correspond to holes in the wall. (b) Numerical solution of Equation (7) at *t* = 50 and *t* = 100. The grey lines indicate the initial condition and the arrow indicates the direction of increasing time. (c) The time evolution of the position of the leading edge of the travelling wave solution. All results are obtained using 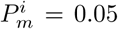, 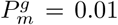, 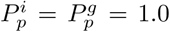, 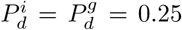, *δx* = 0.1, *δt* = 0.01, *∊* = 10^−6^, *υ* = 0.2760.

Travelling wave behaviour for Equation (7) with positive-negative *F*_*s*_(*C̄*) is shown in Figure 5. The numerical solution of Equation (7), in (*C*, *U*) co-ordinates, passes through the wall of singularities where Equation (10) is finite and forms a heteroclinic orbit between (*S*, 0) and (0, 0). The travelling wave front is of classic type, a result predicted by the analysis performed by Maini *et al*. [38] as *F_s_*(0) ≠ 0 and *F_s_*(1) ≠ 0.

### Allee kinetics

If the birth and death mechanisms are either competitive or co-operative, that is, 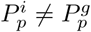 and 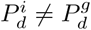, then the source term represents an Allee effect [15] and hence Equation (2) can be expressed as

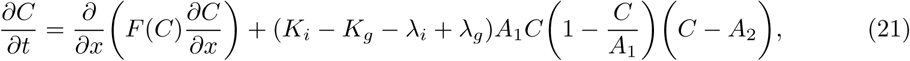

where

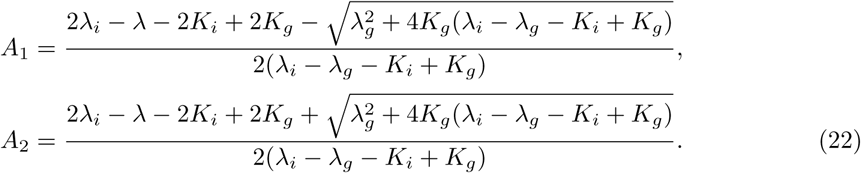

Note that either 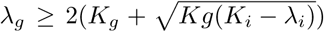 or *λ_i_* > *K_i_* must be satisfied or *R*(*C*) ≤ 0 for 0 ≤ *C* ≤ 1 and the population will tend to extinction. In travelling wave co-ordinates, Equation (21) is

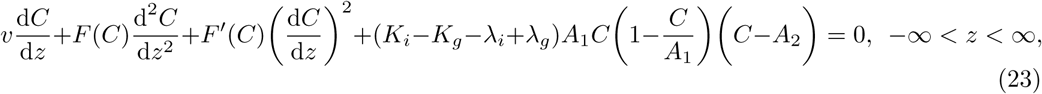

and, making the substitution *U* = d*C*/d*z*, it corresponds to

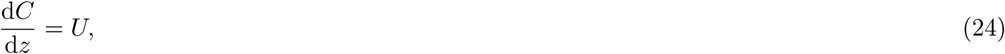

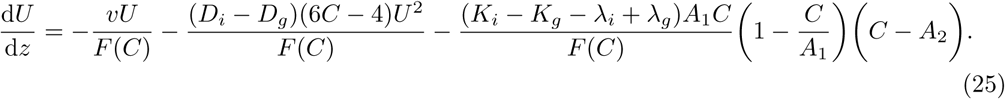

If 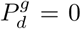, *A*_1_ = 1, and the source term in (21) simplifies to *R*(*C*) = *rC*(1 – *C*)(*C* – A), where *r* = *K_i_* – **λ*_i_* + **λ*_g_* is the intrinsic growth rate and *A* = (*K_i_* – **λ*_i_*)/(*K_i_* – **λ*_i_* + **λ*_g_*) is the Allee parameter [15].

A new variable *C̄* = *C*/*A*_1_ is introduced such that the range of physically relevant *C̄* values corresponds to 0 ≤ *C̄* ≤ 1. Substituting *C̄* into Equation (21) results in

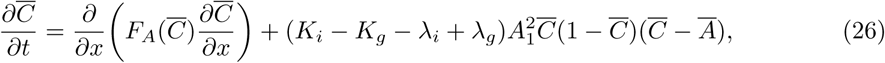

where 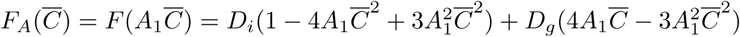 and

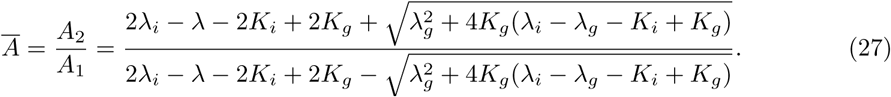

The transformed nonlinear diffusivity, *F_A_*(*C̄*), has the same characteristics as *F*_*s*_(*C̄*), presented in Figure 2, albeit in terms of the scaled Allee carrying capacity, *A*_1_. For 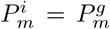, *F_A_*(*C̄*) represents linear diffusion. Reaction-diffusion equations with linear diffusion and either weak or strong Allee kinetics have been well-studied [15, 19, 22, 23, 25, 26, 28, 30]. For additional details we refer the reader to [15]. Weak Allee kinetics correspond to (*K_i_* – *K_g_* – *λ_i_* + *λ_g_*) > 0 and *Ā* < 0, and represent a growth rate that is inhibited at low densities, compared to logistic growth, but remains positive. Strong Allee kinetics correspond to (*K_i_* – *K_g_* – **λ*_i_* + **λ*_g_*) > 0 and 0 < *Ā*< 1 [15], and represent a growth rate that is negative beneath a threshold value, and positive otherwise. Interestingly, a third type of Allee kinetics can arise from the parameter values chosen in the discrete model, that has not been considered previously. If (*K_i_* – *K_g_* – **λ*_i_* + **λ*_g_*) < 0 and *Ā* > 1, the growth rate is non-negative for all relevant *C̄* values but is inhibited at high densities, compared to logistic growth, rather than low densities like the weak Allee effect. We term this type of growth term the reverse Allee effect. Representative source terms showing the three types of Allee effect are compared with a logistic source term in Figure 6.

**Figure 6.**
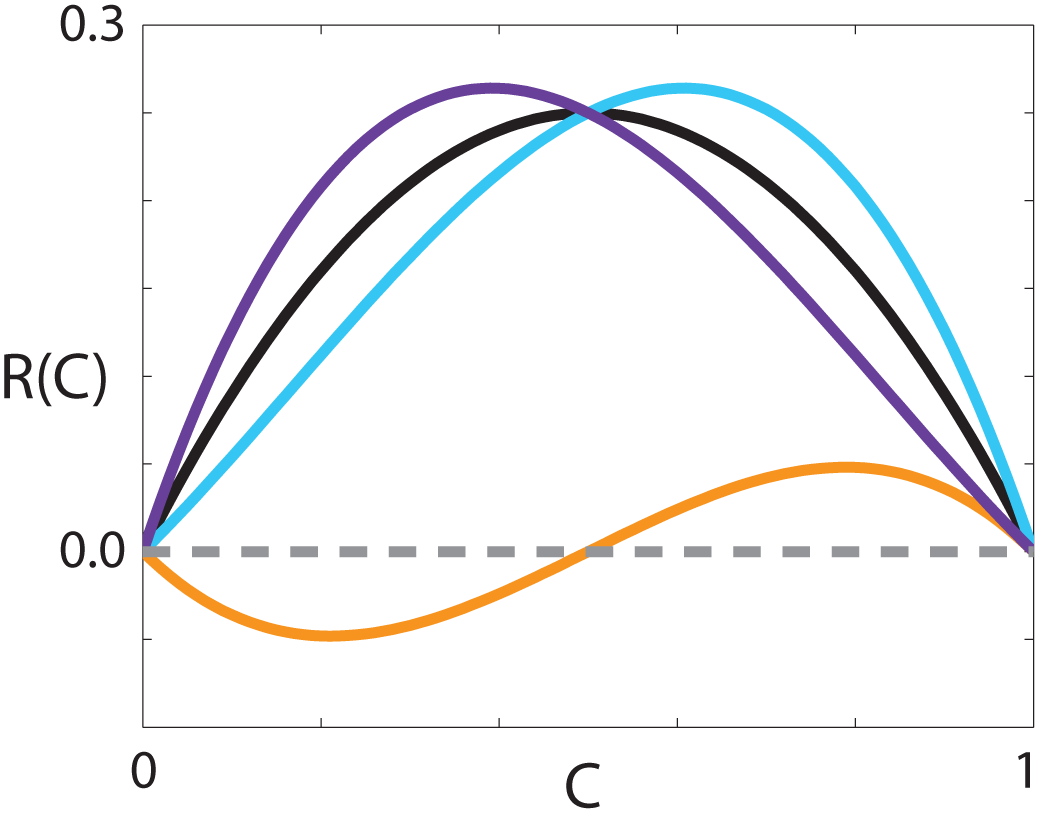
**Comparison of source terms.** *R*(*C*) corresponding to the weak Allee effect with *r* = 1, *A* = −0.5 (cyan), strong Allee effect with *r* = 1, *A* = 0.5 (orange), reverse Allee effect with *r* = −1, *A* = 1.5 (purple) and logistic growth with *r* = 1 (black).

For all combinations of neutral, competitive and co-operative mechanisms that give rise to a reaction-diffusion equation with Allee kinetics we examine the ability of the equation to give rise to long time travelling wave solutions. Furthermore, the three types of Allee effect arising from the discrete model are considered. Several of these cases have been presented and examined previously, but we present details about the travelling wave solutions for all combinations of diffusivity functions and Allee effects in Table 3. A detailed analysis for each case is presented in the Supplementary Material.

**Table 3.**
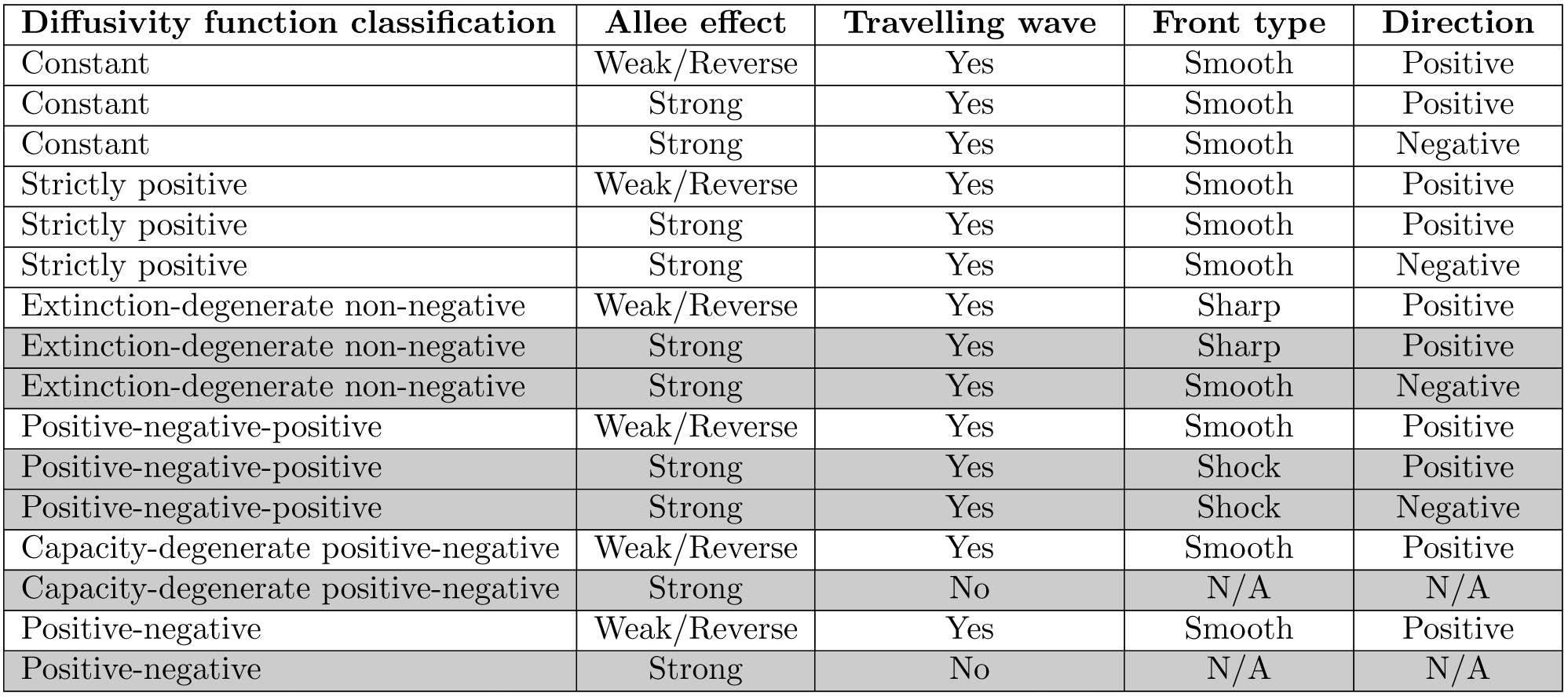
Classification of travelling wave solutions arising from different classes of PDEs with Allee kinetics. Highlighted entries refer to cases analysed in detail in the manuscript; the other cases are analysed in the Supplementary Material.

| Diffusivity function classification | Allee effect | Travelling wave | Front type | Direction |
| --- | --- | --- | --- | --- |
| Constant | Weak/Reverse | Yes | Smooth | Positive |
| Constant | Strong | Yes | Smooth | Positive |
| Constant | Strong | Yes | Smooth | Negative |
| Strictly positive | Weak/Reverse | Yes | Smooth | Positive |
| Strictly positive | Strong | Yes | Smooth | Positive |
| Strictly positive | Strong | Yes | Smooth | Negative |
| Extinction-degenerate non-negative | Weak/Reverse | Yes | Sharp | Positive |
| Extinction-degenerate non-negative | Strong | Yes | Sharp | Positive |
| Extinction-degenerate non-negative | Strong | Yes | Smooth | Negative |
| Positive-negative-positive | Weak/Reverse | Yes | Smooth | Positive |
| Positive-negative-positive | Strong | Yes | Shock | Positive |
| Positive-negative-positive | Strong | Yes | Shock | Negative |
| Capacity-degenerate positive-negative | Weak/Reverse | Yes | Smooth | Positive |
| Capacity-degenerate positive-negative | Strong | No | N/A | N/A |
| Positive-negative | Weak/Reverse | Yes | Smooth | Positive |
| Positive-negative | Strong | No | N/A | N/A |

#### Persistence and extinction

A key question of interest for a particular class of PDE is whether the population described persists or becomes extinct in the long time limit. In all cases with Fisher kinetics with *λ* > *K*, the source term is positive for 0 ≤ *C* ≤ 1, and subsequently the population persists and spreads. As the kinetics representing an Allee effect can contain a source term that is negative for an interval of *C*, it is less obvious whether the minimum wave speed is positive or negative, corresponding to persistence or extinction, respectively.

For the case with constant *F*(*C*) and 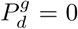, the minimum wave speed for Equation (21) with *A* < −1/2 is 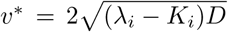 and hence the population persists, provided *λ_i_* > *K_i_*. Introducing 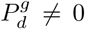 results in the same minimum wave speed, provided that *Ā* < −1/2. This implies that introducing grouped agent death at a rate that does not result in a population tending to extinction has no influence on the invasion speed of the population. Specifically, the condition for *A* < −1/2 with *K_g_* = 0 corresponds to 3(*λ_i_* – *K_i_*) > *λ_g_*. It can be shown that, with 3(*λ_i_* – *K_i_*) > *λ_g_*, we require 3*K_g_* < *λ_g_* for *Ā* < −1/2. This implies that there is a range of *K_g_* values that result in a travelling wave with a minimum wave speed that is independent of both *K_g_* and *λ_g_*. Interestingly, this suggests that if a control is implemented that increases the death rate of grouped agents, there is a threshold value of 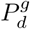 for the control to influence the invasion speed and the subsequent persistence of the population. Introducing a non-zero *K_g_* value for a parameter regime that results in the strong Allee effect with *K_g_* = 0 never changes the type of Allee effect. It is possible to go from a weak Allee effect to a reverse Allee effect by introducing a non-zero *K_g_* value. Non-zero *K_g_* values correspond to a decreased benefit for grouped agents, which explains why the source term, previously a weak Allee effect, becomes the reverse Allee effect, corresponding to inhibited growth at high density.

The reaction-diffusion equation with constant *F_A_*(*C̄*) and the strong Allee effect, corresponding to 0 < *A*_2_ < *A*_1_ ≤ 1, has a unique wave speed 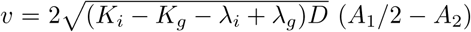 [28]. This implies that for *A*_2_ > *A*_1_/2, *υ* < 0 and *υ* > 0 otherwise. Furthermore, the same wave speed applies for −*A*_1_/2 < *A*_2_ < 0 [28]. For both intervals, the minimum wave speed depends on the *K_g_* value, and hence implementing any kind of partial eradication of the grouped agents will either reduce the speed of invasion or cause the extinction of the population.

For cases where *F_A_*(*C̄*) ≥ 0 for 0 ≤ *C* ≤ 1 and *F_A_*(*C̄*) is not constant, we follow the approach of Hadeler to establish whether the minimum wave speed is positive, and hence the population persists [42–44]. The integral condition for the wave speed to be positive,

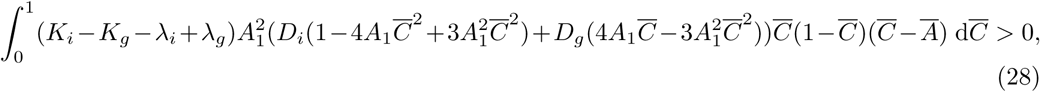

corresponds to

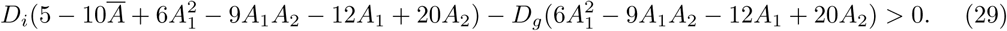

If *D_i_* = *D_g_*, then *Ā* > 1/2 leads to *υ* < 0. For the strong Allee effect, *A*_1_ > *A*_2_ = *ĀA*_1_, we can determine the threshold value for the persistence of the population, namely,

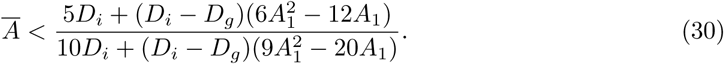

Considering the two limiting cases for strictly positive *F_A_*(*C̄*), where *D_i_* = 0 and *D_i_* = 4*D_g_*, *Ā* takes on a value of 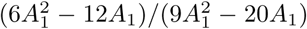 and 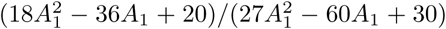, respectively. These values reduce to 6/11 and 2/7 in the case that *A*_1_ = 1, corresponding to *K_g_* = 0. Therefore, populations with isolated agents that are more motile than grouped agents are less susceptible to extinction. To illustrate how the threshold value changes with *A*_1_ 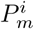 and 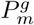, Figure 7 shows the maximum *A*_2_ and *Ā* values for three different 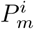 and 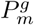 combinations. The *A*_2_ value corresponds to the persistence threshold for a given *A*_1_ value. The *Ā* value can be interpreted as the highest proportion of a given *A*_1_ value that will result in the persistence of the population. For example, in Figure 7(a), we see that with 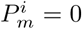 and *A*_1_ = 0.5 we require *A*_2_ < 0.194 for persistence. This corresponds to *Ā* < 0.388.

**Figure 7.**
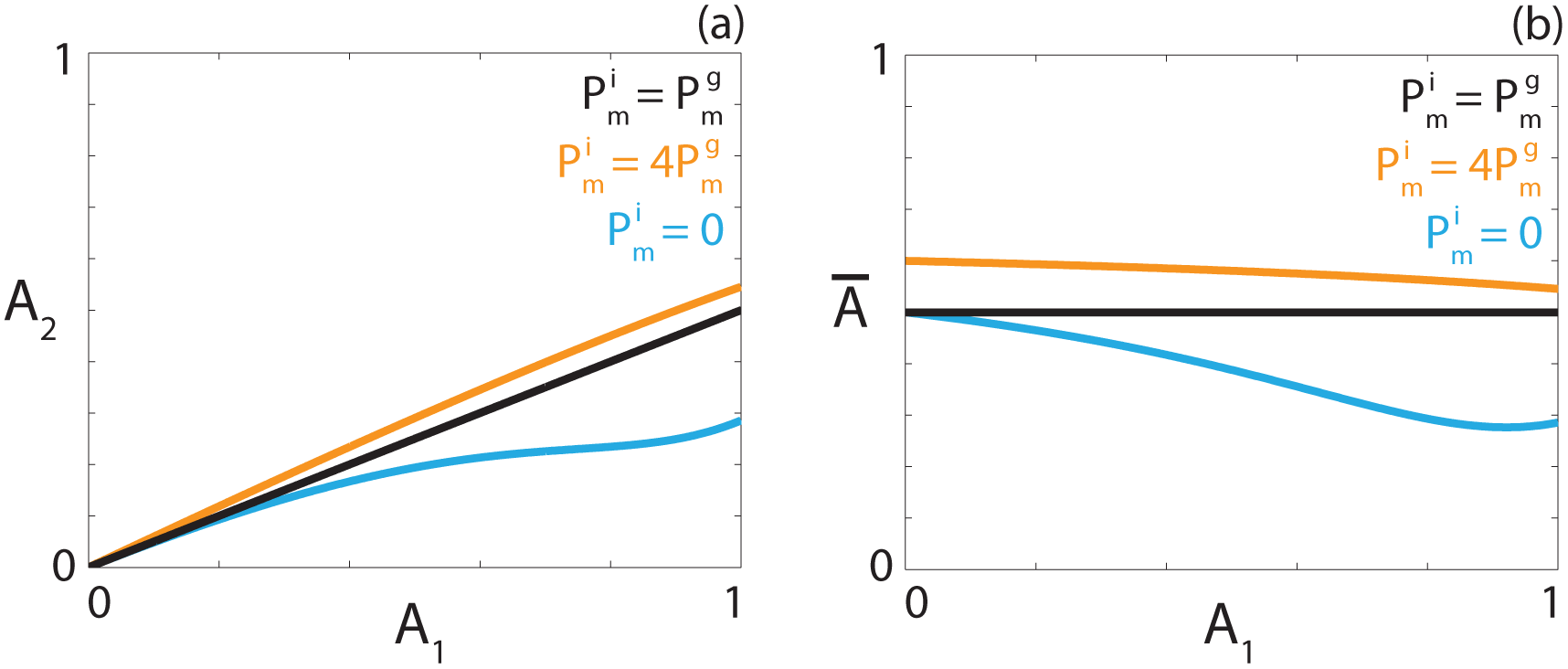
**Persistence threshold.** Persistence threshold as a function of the carrying capacity *A*_1_, expressed as (a) an explicit value; (b) a proportion of the carrying capacity for three different diffusivities, corresponding to 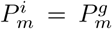 (black), 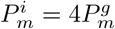 (orange), 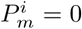 (cyan).

#### Extinction-degenerate non-negative nonlinear diffusivity function

Travelling wave behaviour for the strong Allee effect with extinction-degenerate non-negative *F*(*C*) is shown in Figure 8. The numerical solution of Equation (21) with *A* = 1/4, in Figures 8(a)-(c), leads to a sharp-fronted travelling wave solution near *C* = 0 with *υ* > 0. With *A* = 1/4, we expect to obtain *υ* > 0. For a parameter regime that results in *A* = 4/7, we obtain a travelling wave solution of Equation (21) with *υ* < 0 (Figures 8(d)-(f)). Interestingly, the sharp front near *C* = 0 is not present for the strong Allee effect with *υ* < 0.

**Figure 8.**
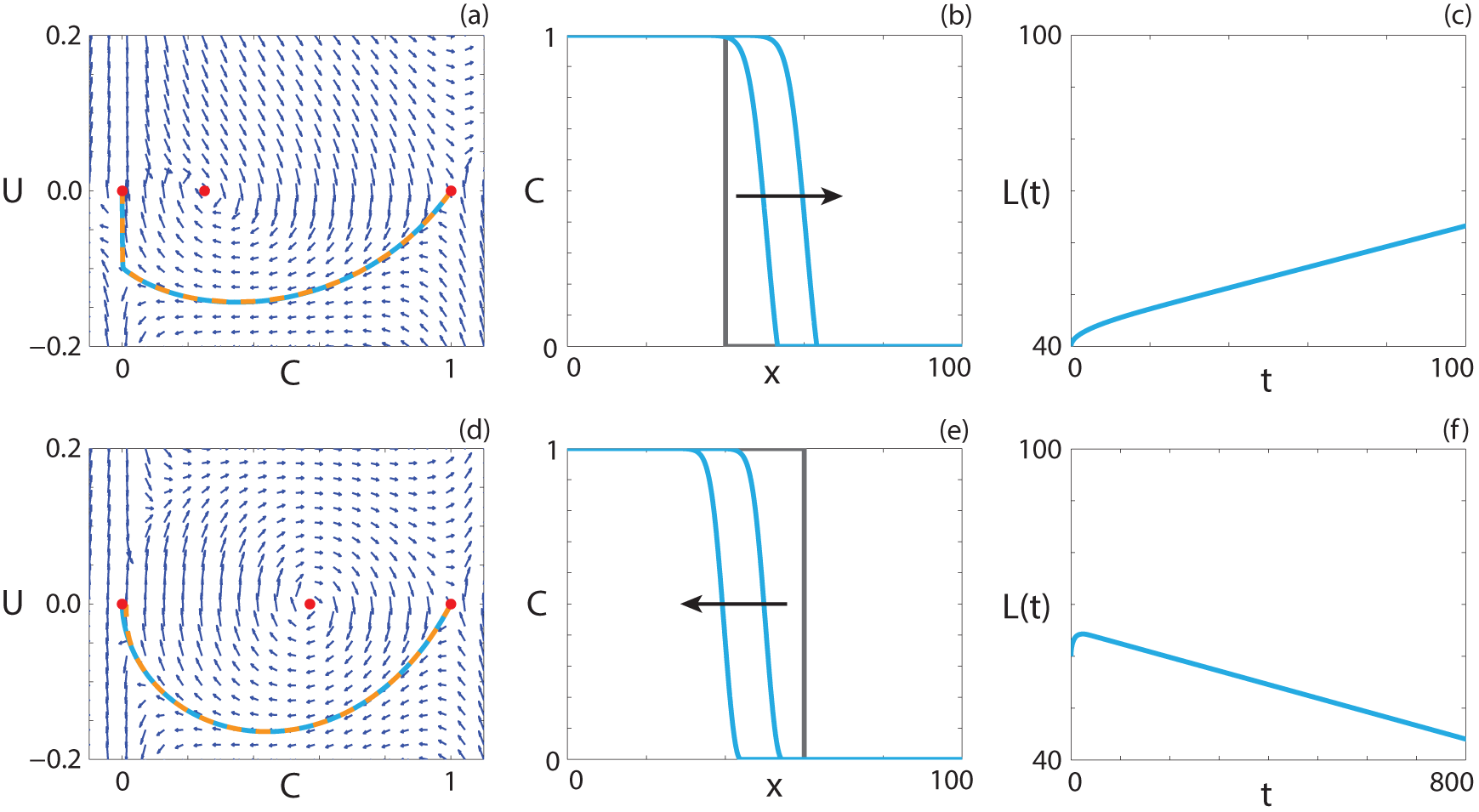
**Travelling wave behaviour for Equation (21) with the strong Allee effect and extinction-degenerate non-negative *F*(*C*) (Case 6.2).** (a), (d) Phase plane for the system (24)–(25) with the numerical solution of Equations (21) (cyan, solid) and (23) (orange, dashed), in (*C*, *U*) co-ordinates, superimposed. Red circles correspond to equilibrium points. (b), (e) Numerical solution of Equation (21) calculated at (b) *t* = 50 and *t* = 100; (e) *t* = 400 and *t* = 800. The grey lines indicate the initial condition and the arrows indicate the direction of increasing time. (c), (f) The time evolution of *L*(*t*). All results are obtained with *δx* = 0.01, *δt* = 0.005, *∊* = 10^−6^, 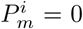, 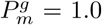, 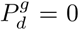, (a)-(c) 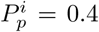, 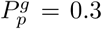, 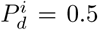, *υ* = 0.199; (d)-(f) 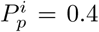, 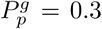, 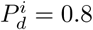, *υ* = −0.026.

#### Positive-negative-positive nonlinear diffusivity function

A positive-negative-positive *F*(*C*), where there is an interval *α* < C < *β* where *F*(*C*) < 0, corresponds to parameter pairs highlighted in purple in Figure 2(a). Kuzmin and Ruggerini [37] examine reaction-diffusion equations with similar properties for the strong Allee effect, in the context of diffusion-aggregation models, and provide conditions for smooth travelling wave solutions to exist. For a solution with *υ* > 0, we require *A* < *α* [37] and

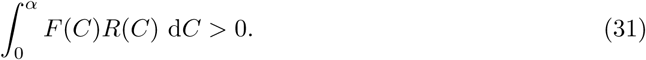

Furthermore, we require [37]

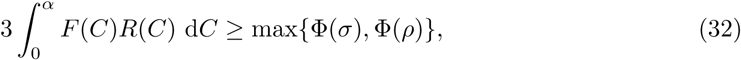

**Figure 9.**
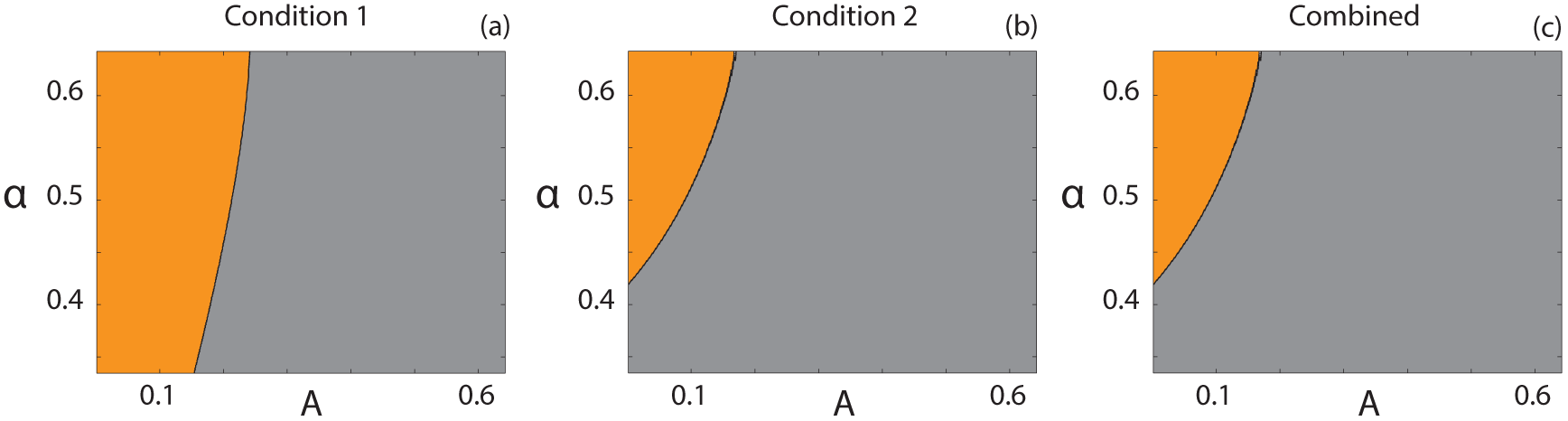
**Parameter pairs that satisfy Kuzmin and Ruggerini's Conditions.** (a) Condition (31); (b) Condition (32); (c) Conditions (31)–(32) combined. Orange regions correspond to parameter pairs that satisfy the respective condition(s), whereas grey regions correspond to parameter pairs that do not.

where 

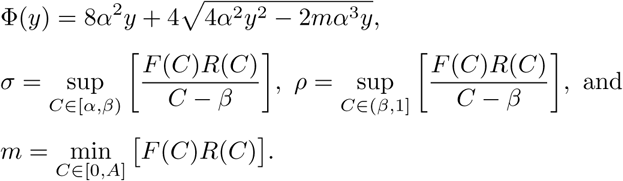

A suite of 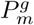 values with 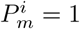 that correspond to 1/3 < *α* < 2/3 are considered for parameter regimes that result in *A* < *α*. Figures 9(a)-(c) show the parameter pairs, (*A*, *α*), that satisfy Condition (31), Condition (32) and Conditions (31)–(32) simultaneously, respectively. Orange regions represent parameter pairs where the condition is satisfied and grey regions represent parameter pairs where the condition is not satisfied. These results suggest that smooth travelling wave solutions should exist for certain choices of parameters. Interestingly, all parameter pairs that satisfy Condition (31) also satisfy Condition (32).

For Equation (7) with positive-negative-positive *F*(*C*), smooth travelling wave solutions that pass through holes in the wall of singularities for positive-negative-positive *F*(*C*) are obtained. The minimum wave speed bound presented by Ferracuti et al. [36] implies that the locations of the holes in the wall occur are real-valued for the wave speed arising from the Heaviside initial condition. As such, to obtain smooth travelling wave solutions of Equation (21) with positive-negative-positive *F*(*C*), we might expect that the wave speed satisfies 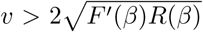, such that the holes in the wall at *C* = *β* are real-valued.

**Figure 10.**
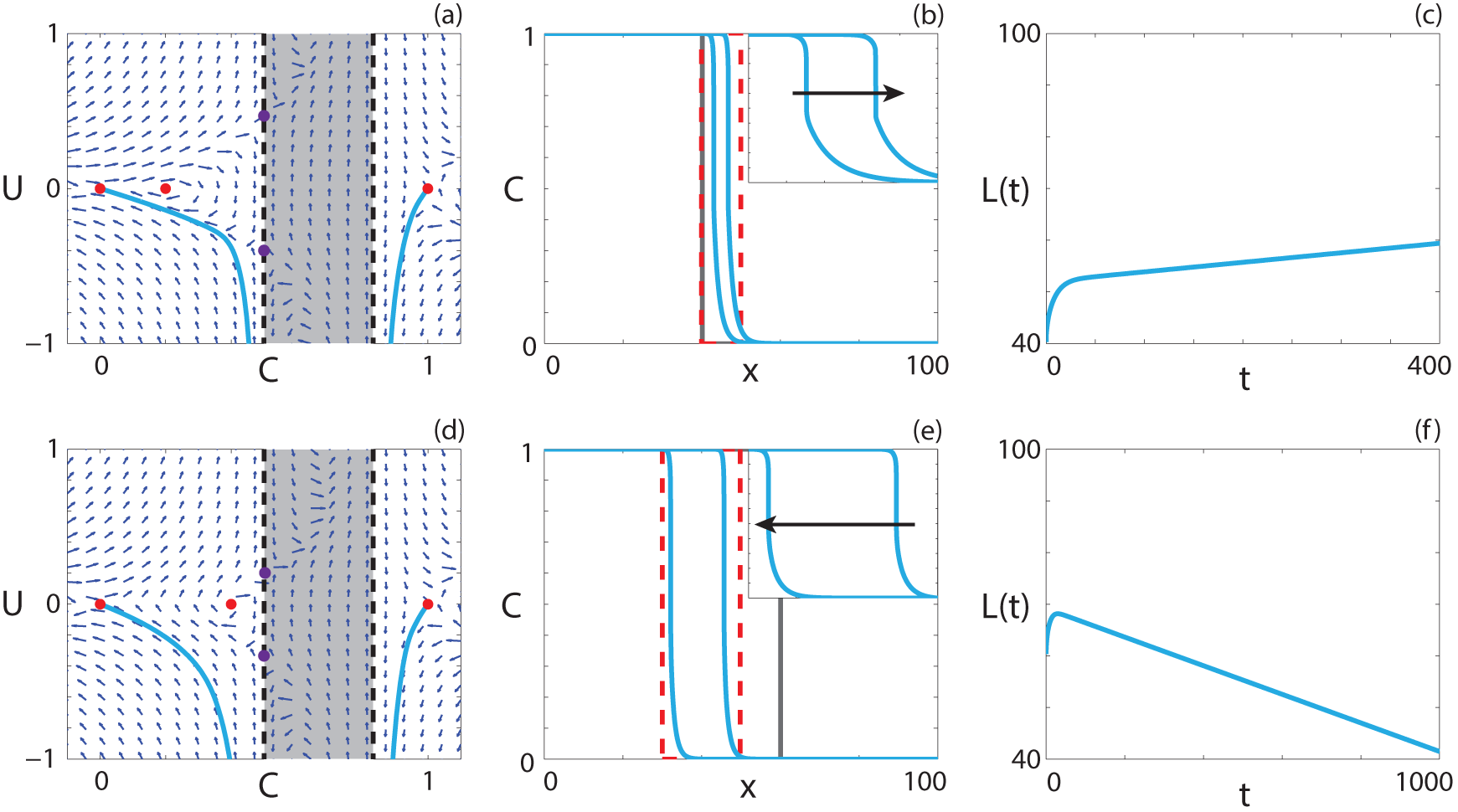
**Travelling wave behaviour for Equation (21) with the strong Allee effect and positive-negative-positive *F*(*C*) (Case 6.3).** (a), (d) Phase plane for the system (24)–(25) with the numerical solution of Equation (21) (cyan, solid), in (*C*, *U*) co-ordinates, superimposed. The dashed black lines denote a wall of singularities. Red circles correspond to equilibrium points and purple circles correspond to holes in the wall. (b), (e) Numerical solution of Equation (21) calculated at (b) *t* = 200 and *t* = 400; (e) *t* = 500 and *t* = 1000. The grey lines indicate the initial condition and the arrows indicate the direction of increasing time. The insets correspond to the areas within the red dashed lines, and highlight the shocks. (c), (f) The time evolution of *L*(*t*). All results are obtained with *δx* = 0.05, *δt* = 0.001, *∊* = 10^−6^, 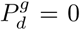, (a)-(c) 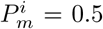, 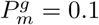, 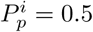, 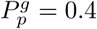, 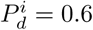, *υ* = 0.009; (d)-(f) 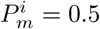, 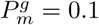, 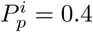, 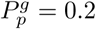, 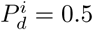, *υ* = −0.028.

Following the approach used for Equation (7) with positive-negative-positive *F*(*C*), it is simple to demonstrate that both the weak and reverse Allee effect have real-valued holes in the wall (Supplementary Material). We now examine numerical solutions of Equation (21) with the strong Allee effect. For parameter regimes that give rise to wave speeds that satisfy 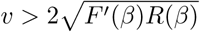, numerical travelling wave solutions could not be found. While the condition for real-valued holes in the wall is satisfied, the zeroes of Equation (25) are imaginary for a certain interval of *C > β*. This corresponds to a nullcline that is not real-valued for certain *C* values.

We now consider parameter regimes corresponding to the strong Allee effect with the additional restriction that 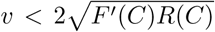 for 2/3 < *C* ≤ 1. For all 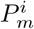 and 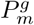 that give rise to a positive-negative-positive *F*(*C*), holes in the wall at *C* = *β* do not exist and, as such, we do not expect to obtain smooth solutions. Interestingly, we observe travelling wave solutions with shocks such that the solution never enters the region *α* < *C* < *β*. An example of a shock-fronted travelling wave solution for the strong Allee effect with both *υ* > 0 and *υ* < 0 is shown in Figures 10(a)-(c) and Figures 10(d)-(f), respectively. Solutions of diffusion equations, without any source terms, that contain shocks have been reported previously [50, 51]. Similarly, shock-fronted travelling wave solutions arise in other kinds of models, including multispecies models of combustion [54] and haptotactic cell migration [53]. However, the models presented here are very different as our model contains a source term and no advection term, and it is therefore of interest to determine the properties of the reaction-diffusion equation that lead to shock-fronted travelling wave solutions.

#### Capacity-degenerate positive-negative nonlinear diffusivity function

Capacity-degenerate positive-negative *F*(*C*), where *F*(1) = 0, arises if 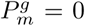 and includes an interval 1/3 < *C* < 1 where *F*(*C*) < 0. For the corresponding case with Fisher kinetics, despite the degenerate nature of the nonlinear diffusivity function at *C* = 1, we did not obtain solutions with a sharp front near *C* = 1. Instead, the solution passes through the region of negative diffusivity and a hole in the wall at *C* = 1/3, leading to smooth travelling wave solutions. As such, we expect similar solutions for both the weak and reverse Allee effect due to the qualitatively similar behaviour of the *R*(*C*) function. It is of interest to examine whether smooth or shock-fronted travelling wave solutions arise from Equation (21) for the strong Allee effect and capacity-degenerate positive-negative *F*(*C*), as for the positive-negative-positive *F*(*C*) no smooth travelling wave solutions could be found.

As expected, smooth travelling wave solutions for both the weak and reverse Allee effects with capacity-degenerate positive-negative *F*(*C*) are obtained. The solution behaviour for both the weak and reverse Allee effects are presented in the Supplementary Material. For the strong Allee effect, we examined a considerable number of parameter regimes and initial conditions and were unable to find travelling wave solutions.

#### Positive-negative nonlinear diffusivity function

For the case where *F_A_*(*C̄*) has exactly one zero on the interval 0 ≤ *C* ≤ 1 at *C* = *ω*, Maini *et al*. [39] examine the existence of travelling wave solutions, and provide the necessary conditions for existence,

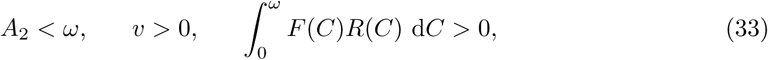

where *F*(*ω*) = 0 and 0 < *ω* < 1. For the strong Allee effect in this parameter regime, the third part of Condition (33) corresponds to

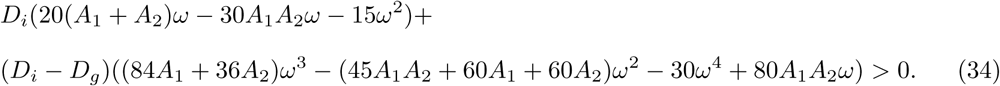

Equation (21) is equivalent to

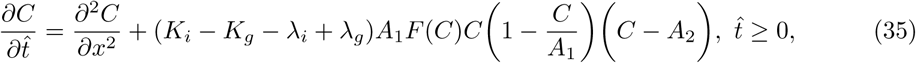

on the interval 0 ≤ *C* < *ω*, and equivalent to

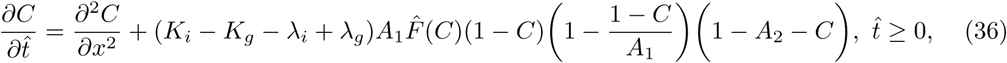

where *F̂*(*C*) = −*F*(1 – *C*), on the interval *ω* ≤ *C* ≤ *A*_1_. The final necessary and sufficient condition from Maini *et al*. [39] for the existence of travelling wave solutions is that the minimum wave speed for Equation (35), 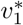, is greater than, or equal to, the minimum wave speed for Equation (36), 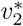. On the interval 0 ≤ *C* ≤ *ω*, Equation (21) has a strictly positive *F*(*C*), where *F*(*C*) ≤ *D_i_*, and strong Allee kinetics. Hence, the minimum wave speed for Equation (35) has an upper bound, 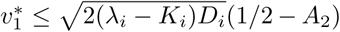. On the interval *ω* < *C* < *A*_1_ Equation (36) has a source term qualitatively similar to the Fisher-Kolmogorov equation and hence a lower bound for the minimum wave speed exists [39], 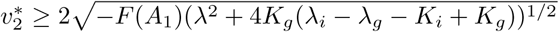. For all parameter regimes considered that correspond to the strong Allee effect with positive-negative *F_A_*(*C̄*) we never observe a case where the upper bound for 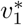 is higher than the lower bound for 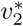 and hence the conditions required for travelling wave solutions are not met. As expected, numerical solutions of Equation (21) in these parameter regimes did not lead to travelling wave behaviour.

## Discussion

In this work we present a discrete lattice-based model of birth, death and movement. The model is an exclusion process, and hence it explicitly incorporates crowding by allowing no more than one agent per site. A key feature of the model is that birth, death and movement rates depend on whether an agent is isolated or whether it is part of a group of agents. The discrete model can, therefore, be used to describe co-operative or competitive mechanisms [8, 13–15]. These kinds of mechanisms are thought to be relevant to many applications in cell biology [8,12,55,56] and ecology [13–15]. By considering different combinations of parameters, the continuum limit PDE approximation of the discrete model leads to 22 different cases. These cases are reaction-diffusion equations with either Fisher kinetics or Allee kinetics, and a variety of density-dependent nonlinear diffusivity functions (Table 1). This approach also leads to a new kind of Allee effect, which we call the reverse Allee effect, where the growth rate is inhibited at high density. Although some of the PDEs that we consider have been investigated previously [15, 18–30, 36–41], they have never been linked together before using a single modelling framework. The presence of Allee kinetics allows for the more realistic description of biological and ecological phenomena, as the standard reaction-diffusion model with Fisher kinetics predicts either the population tending to extinction everywhere or the spread of the population in the form of a travelling wave. In comparison, Allee kinetics can describe population retreat, as well as shocks in the invading front of a population. Well-defined edges are thought to be present in invasive tumours [56], which can be described with travelling waves containing shocks.

In this work, we summarise properties of the long time travelling wave solutions for all classes of PDEs arising from our discrete model. For certain PDEs, where only existence of travelling wave solutions has been considered previously, we present numerical solutions here for the first time. We find that PDE models with density-dependent nonlinear diffusivity functions that have regions of negative diffusivity require a sufficiently non-negative source term to support smooth travelling wave solutions. Furthermore, there appears to be a threshold proliferation value, depending on the rate of motility, that must be exceeded for travelling wave solutions to be observed numerically. However, we do not comment on the putative relationship between the parameters in the discrete model and the existence of travelling wave solutions in the continuum limit PDE. Interestingly, for the strong Allee effect, shock-fronted travelling wave solutions are obtained. Following arguments presented in [39], we show that smooth travelling wave solutions cannot be obtained for certain types of nonlinear diffusivity functions and the strong Allee effect. We describe how nonlinear diffusion can either hinder or promote the persistence of a population, depending on the relative motility rates of the isolated and grouped agents. Interestingly, the motility rates affect the persistence differently for different carrying capacities. This relationship could provide insight into the requirements for a cell population, for example, to persist in the presence of a chemical treatment.

The six birth, death and motility rate parameters in the discrete model allow for the interpretation of the results in terms of whether individuals are part of, or isolated from, the bulk population. For example, a parameter regime corresponding to the strong Allee effect with constant diffusivity and no grouped agent death leads to the same travelling wave speed in the PDE description as a parameter regime corresponding to the strong Allee effect with constant diffusivity and a non-zero rate of grouped agent death, up to a threshold. This implies that a sufficiently strong intervention strategy aimed at grouped agents must be implemented if the goal of the intervention is to slow or halt the invasion of a population.

The work presented here suggests several avenues for future research. This work could be generalised by considering a two- or three–dimensional discrete process and deriving the continuum limit PDE descriptions in higher dimensions. This kind of higher-dimensional model might provide a more accurate description of real world observations where one-dimensional travelling wave solutions might not apply. In this work, numerical travelling wave solutions for each class of PDE are examined, but the formal stability of these travelling wave solutions is not considered. Another approach for analysing the discrete model would be to consider a coupled multispecies PDE model by accounting for the density of isolated agents and the density of grouped agents separately. This approach would lead to a system of two coupled PDEs instead of a single PDE for the total agent density. However, instead of working with coupled multispecies PDEs, we have taken the simplest and most fundamental approach of considering a single PDE description of the total population. In addition, a significant number of mechanisms could be implemented into the discrete model, such as cell-to-cell adhesion/repulsion [57, 58] or directed migration of isolated agents, such as chemotaxis [59]. We leave these extensions for future analysis.

## Methods

### Discrete model

We consider a one-dimensional lattice-based random walk with *X* sites and lattice spacing ∆ [60]. Each site may be occupied by, at most, one agent [61–63]. The number of agents at time *t* is *N*(*t*). Agents attempt to undergo birth, death and movement events. During a birth event, an agent attempts to place a daughter agent at a randomly selected nearest-neighbour site. This event is successful provided that the selected site is vacant. During a death event, an agent is removed from the lattice. During a movement event, an agent attempts to move to a randomly selected nearest-neighbour site. This event is successful provided that the selected site is vacant. We distinguish between types of agents based on the number of occupied nearest-neighbour sites for each agent [64]. We refer to agents with zero occupied nearest-neighbour sites as *isolated agents*, and agents with one or two occupied nearest-neighbour sites as *grouped agents*. This approach allows us to specify different birth, death and movement rates for isolated and grouped agents.

Different parameter choices can be used to impose either co-operative or competitive mechanisms, where an increase in local agent density provides a positive or negative benefit, respectively. Specifically, in situations where the group motility or group proliferation rates are higher than the isolated motility or isolated proliferation rates, respectively, we interpret this choice of parameters as a model of co-operation. Similarly, in situations where the group motility or group proliferation rates are lower than the isolated motility or isolated proliferation rates, respectively, we interpret this as a model of competition.

During each time step of duration *τ*, *N*(*t*) agents are selected at random, one at a time, with replacement, and are given the opportunity to undergo a movement event. The constant probability that a selected agent attempts to undergo a movement event is 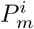 for an isolated agent and 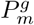 for a grouped agent. We repeat this process for both birth and death events, with respective constant probabilities 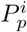 and 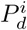 for isolated agents and 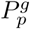 and 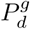 for an agent within a group. At the end of each time step we update *N*(*t* + *τ*). To obtain the average agent density at each lattice site we perform *M* identically-prepared realisations of the discrete model and average the binary lattice occupancy at each lattice site at each time step. In any single realisation of the discrete model we have *C_j_* = 1 when site *j* is occupied and *C_j_* = 0 when site *j* is vacant. To evaluate the average occupancy of any lattice site we consider an ensemble of *M* identically-prepared realisations and calculate 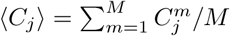.

### Numerical techniques

Here we describe the techniques used to obtain numerical solutions of Equation (2), the corresponding ODE in travelling wave co-ordinates, and to generate the phase planes in (*C*, *U*) co-ordinates.

#### Partial differential equations

To obtain numerical solutions of Equation (2), we first spatially discretise Equation (2) onto a grid with uniform grid spacing *δx* by approximating the spatial derivatives with a central finite difference approximation. A backward Euler approximation with constant time steps of duration *δt* is used to approximate the temporal derivative. The resulting system of nonlinear algebraic equations is solved using Picard iteration with absolute convergence tolerance *∊*. The resulting system of tridiagonal algebraic equations is solved using the Thomas algorithm [68]. All results presented correspond to sufficiently small choices of *δx*, *δt* and *∊* so that the numerical solutions are grid independent. In all cases consider zero-flux boundary conditions are considered, and the finite domain is sufficiently large such that the numerical solution of Equation (2) does not interact with the boundaries on the time scale of the numerical simulations. All numerical solutions correspond to a Heaviside initial condition with *C* = 1 for *x* ≤ *X*_0_, and *C* = 0 otherwise.

#### Ordinary differential equations

The second order ODEs in the travelling wave co-ordinates are solved using Matlab's `ode45` routine [69]. This routine implements an adaptive Runge-Kutta method with relative error tolerance of 10^−3^ and an absolute error tolerance of 10^−6^ [69]. Travelling wave ODEs that contain a singularity are not solved numerically. Therefore, for these singular problems we obtain only the numerical solution of the PDE and present this solution in the transformed (*C*, *U*) travelling wave co-ordinate system.

#### Phase planes

To generate phase planes we substitute *U* = d*C*/d*z* into the second order travelling wave ODE to obtain a system of two first-order ODEs. The phase plane is constructed by considering 22 equally-spaced values of *C* and 22 equally spaced values of *U* to calculate both d*C*/d*z* and d*U*/d*z* at all 22 × 22 = 484 pairs of (*C*, *U*) values. In each phase plane the same 22 equally spaced values of *C* on the interval 0 ≤ *C* ≤ 1 are considered. However, depending on the steepness of the waveform, we choose a different interval of *U* to construct the phase plane, and this choice is made to accommodate the heteroclinic orbit. The phase planes are constructed using Matlab's `quiver` function. The location of the equilibrium points, where d*C*/d*z* = d*U*/d*z* = 0 are superimposed. Furthermore, in many cases the expression for d*U*/d*z* has a rational form, d*U*/d*z* = *G*(*C*, *U*)/*H*(*C*, *U*). In these cases both the wall of singularities (*H*(*C*, *U*) = 0) and the locations of the holes in the wall (*H*(*C*, *U*) = *G*(*U*, *C*) = 0) are also superimposed.

## Acknowledgements

This work is supported by the Australian Research Council (DP140100249, FT130100148). We thank the referees for their helpful comments.

## Author contributions

STJ, REB, DLSM and MJS conceived the experiments, STJ performed the experiments, STJ, REB, DLSM and MJS analysed the results, STJ and MJS wrote the manuscript. All authors read and approved the final version of the manuscript.

## Additional information

**Competing financial interests**. The authors declare no competing financial interests.

